# Kaptin controls neuronal circuit assembly by limiting actin-driven axon collateral branching

**DOI:** 10.64898/2026.07.27.740990

**Authors:** Roopsali Banerjee, Suneha Mukherjee, Aurnab Ghose

## Abstract

Axon collateral branching is a fundamental determinant of neuronal connectivity, enabling individual neurons to innervate multiple targets and establish functional neural circuits. Although branch initiation requires local actin remodelling within the axon shaft, the mechanisms that restrict actin assembly to prevent excessive branching remain poorly understood. Here, we identify Kaptin (Kptn) as a conserved negative regulator of axon collateral formation that limits the maturation of axonal actin patches. Using primary neuronal cultures, quantitative live-cell imaging, and zebrafish genetics, we show that loss of Kptn enhances the conversion of actin patches into filopodial protrusions, leading to excessive collateral branching. Kptn-deficient zebrafish exhibit increased motor axon arborisation, elevated neuromuscular junction density and impaired motor behaviour. Mechanistically, Kptn antagonises the actin elongation factor Formin-2 (Fmn2) to regulate actin filament barbed-end dynamics, thereby controlling the threshold for productive branch initiation. Importantly, these branching defects occur independently of Kptn’s established role in mTORC1 signalling, revealing a distinct physiological function during neuronal development. Our findings identify Kptn as a key molecular brake that constrains axon collateral branching and establish negative regulation of actin patch maturation as a fundamental mechanism controlling neuronal circuit assembly. This work provides a mechanistic framework for understanding how KPTN mutations associated with intellectual disability and epilepsy disrupt neuronal connectivity.

## INTRODUCTION

Axonal branching is a central feature of neuronal morphogenesis that determines the architecture and topology of neural circuits by regulating the number and distribution of synaptic connections across target populations (Gibson and Ma, 2011; Kalil and Dent, 2014; Ziak et al., 2026). Through axon collateral branching—the de novo formation of interstitial branches from a pre-existing axon shaft—single neurons innervate multiple targets, thereby increasing the computational capacity and connectivity of neural circuits (Bodakuntla et al., 2021; Gibson and Ma, 2011). Because the density and spatial distribution of axon collaterals critically influence circuit function and behavioural output, the formation of collateral branches must be tightly regulated (Armijo-Weingart and Gallo, 2017).

Accordingly, perturbations in axonal branching are associated with a broad spectrum of neurodevelopmental and neuropsychiatric disorders (Kalil and Dent, 2014; McFadden and Minshew, 2013; Van Battum et al., 2015). Excessive axonal branching can disrupt synaptic scaling and promote network hyperconnectivity, a feature observed in subsets of autism spectrum disorders and epileptic encephalopathies (McFadden and Minshew, 2013). Conversely, reduced or aberrant branching compromises target innervation and circuit assembly, contributing to intellectual disability, developmental delay and motor dysfunction (Ramakers, 2002).

The architecture of neuronal arbourisations is constrained by energetic and material costs, requiring a balance between computational efficiency and resource allocation (Chklovskii et al., 2002; Ramón y Cajal, 1995). Interstitial axonal branching near target fields has been proposed to represent a developmental implementation of geometric optimisation principles that minimise wiring costs (Cherniak, 1992). However, although such principles specify favourable branching geometries, the initiation of collateral branches is governed by local cytoskeletal remodelling within the axon shaft. Defining the cytoskeletal mechanisms that regulate collateral branch initiation, therefore, remains central to understanding the relationship between neuronal structure and function.

Axon collateral formation is initiated by extracellular cues, including nerve growth factor (NGF), which induce the formation of axonal F-actin patches through local recruitment of the WAVE1-Arp2/3 machinery (Ketschek and Gallo, 2010). The maturation of these F-actin structures into axonal protrusions requires precise regulation of actin filament barbed-end dynamics, balancing filament elongation with mechanisms that restrain actin assembly (Armijo-Weingart and Gallo, 2017; Gallo, 2024; Kalil and Dent, 2014). Positive regulators of F-actin elongation, including Ena/VASP proteins and the formin Fmn2, promote actin patch maturation and branch initiation (Dwivedy et al., 2007; Kundu et al., 2022; Lebrand et al., 2004; Nagar et al., 2021). However, despite the central role of barbed-end dynamics in controlling axonal branching, the molecular mechanisms that constrain actin patch maturation and prevent excessive collateral formation remain poorly understood.

Kaptin (encoded by the KPTN gene) has recently emerged as a candidate regulator of this negative control network. Kaptin (Kptn) localises to diverse actin-rich structures, including cochlear stereocilia, fibroblast lamellipodia, neuronal growth cones and synapses (Baple et al., 2014; Bearer and Abraham, 1999). Loss-of-function mutations in KPTN are associated with a neurodevelopmental syndrome characterised by intellectual disability, macrocephaly and epilepsy (Baple et al., 2014; Biswas et al., 2025; Horn et al., 2022; Pajusalu et al., 2015; Rawlins et al., 2026; Thiffault et al., 2020; Ullah et al., 2022). Recent biochemical studies demonstrated that Kptn binds directly to the barbed end of F-actin and suppresses filament elongation, establishing it as a direct negative regulator of actin assembly (Dutta et al., 2026). These findings raise the possibility that Kptn constrains axonal branching by limiting the maturation and protrusive potential of axonal actin patches.

Here, using primary neuronal cultures and zebrafish genetics, we identify Kptn as a key actin-remodelling factor that constrains axonal collateral branching. We show that Kptn limits the maturation of axonal actin patches by reducing their probability of transitioning into filopodial protrusions and that this inhibitory activity is functionally opposed by the actin elongation factor Fmn2. Consistent with these findings, loss of either Kptn or Fmn2 disrupts collateral branch formation in zebrafish motor neurons and impairs motor behaviour. Although Kptn has also been identified as a component of the lysosomal KICSTOR complex that suppresses mTORC1 signalling (Wolfson et al., 2017), we demonstrate that its actin remodelling activity, rather than its mTOR-regulatory function, is required for the establishment of normal axonal branching patterns. Together, our findings identify Kptn as a critical negative regulator of actin-driven axonal morphogenesis and establish direct regulation of actin barbed-end dynamics as a key mechanism governing neuronal circuit assembly and function.

## RESULTS

### 1. Kptn null zebrafish exhibit deficits in motor responses

Kaptin (Kptn) is well conserved within the vertebrate lineage with 63.18% sequence identity between the zebrafish and human orthologs (Figure S1 and (Dutta et al., 2026). Whole-mount mRNA *in situ* hybridisation in developing zebrafish embryos revealed expression of *kptn* throughout early development, starting from the one-cell stage (indicating maternal deposition). *kptn* mRNA is detected ubiquitously, but from 24 hours post fertilisation (hpf) it became enriched in the forebrain, midbrain, hindbrain, and spinal cord. (Figure S2).

To investigate Kptn’s function in the nervous system, we generated two independent homozygous *kptn* mutant lines carrying 5 bp and 14 bp deletions in exon 1. Each deletion introduces a frameshift predicted to yield a truncated protein of 43 and 46 amino acids (full length protein is 436 amino acids long), respectively (Figure S3). All subsequent experiments were performed on a heteroallelic line generated by crossing the two homozygous mutant lines (*kptn^Δ5/ Δ14^*).

Morphometric analysis of 3 dpf zebrafish revealed no significant difference between *kptn^+/+^* (3.03 ± 0.02 mm) and *kptn^Δ5/ Δ14^* (2.97 ± 0.03 mm) for body length. Similarly, no difference was observed for the head area between *kptn^+/+^* (0.22 ± 0.01 mm^2^) and *kptn^Δ5/ Δ14^* (0.22 ± 0.01 mm^2^) and eye-to-eye distance for *kptn^Δ5/ Δ14^* (0.46 ± 0.01 mm) with respect to *kptn^+/+^* (0.46 ± 0.01 mm) (Figure S4).

Spontaneous Tail Coiling (STC; a sensory-independent spontaneous motor behaviour in early embryos while within the chorion) and Touch Evoked Escape Response (TEER; a stimulus-evoked flight response) were evaluated in *kptn* null animals, as these represent the earliest coordinated behaviours observable during development (Kohashi and Oda, 2008; Saint-Amant and Drapeau, 1998). Examination of *kptn^Δ5/ Δ14^* at 24 hpf revealed an increased frequency of tail twitching (6.91 ± 0.23 mins^-1^) relative to wild-type (*kptn^+/+^*) zebrafish (4.95 ± 0.21 mins^-1^). Furthermore, *kptn^Δ5/ Δ14^* (8.14 ± 0.13 a.u.) displayed significantly decreased amplitude of the tail twitches compared to the *kptn^+/+^* animals (10.24 ± 0.24 a.u.) (Figure 1A-C).

**Figure 1.**
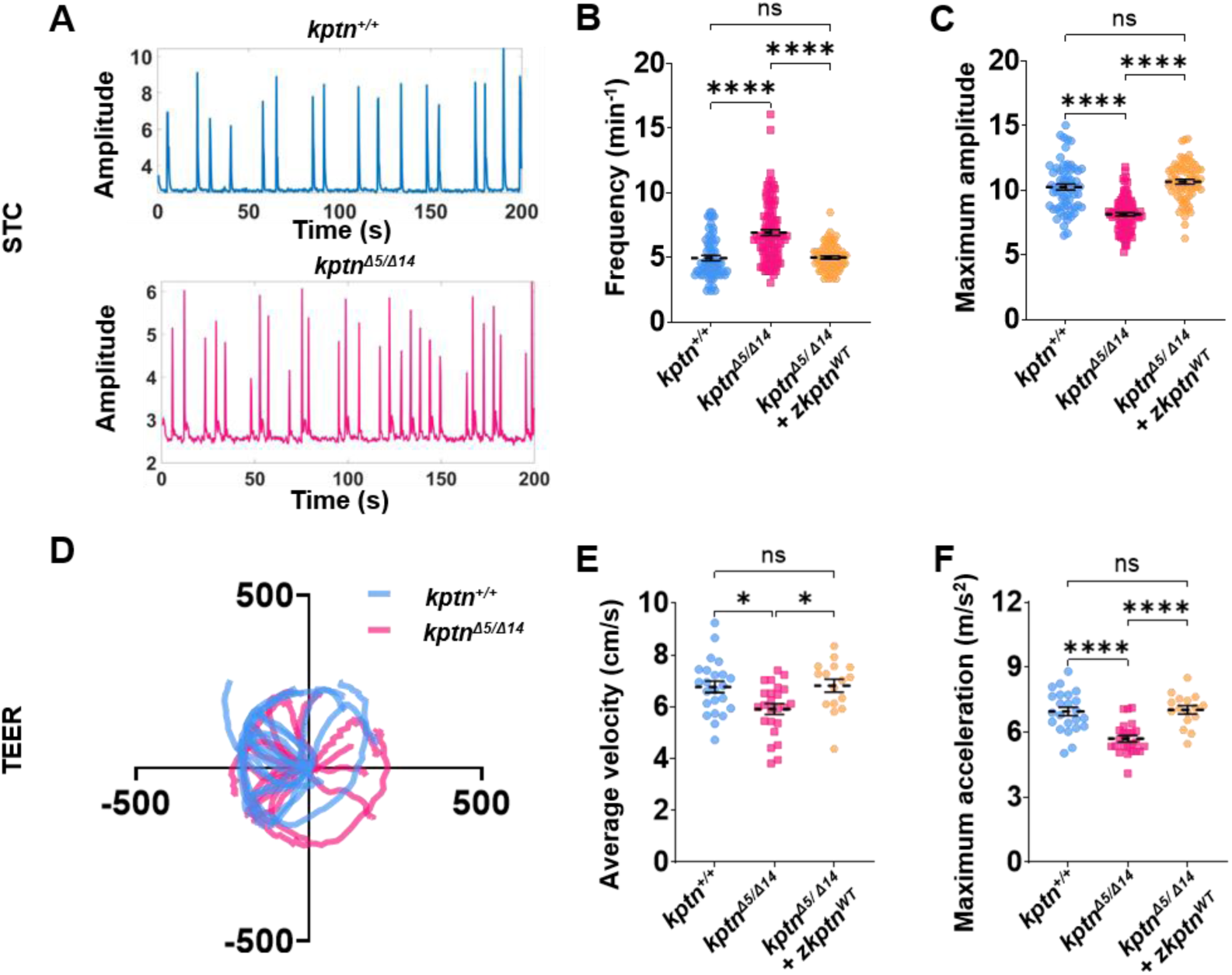
*kptn* mutants exhibit deficits in motor behaviours. (A) Representative plots for *kptn^+/+^* and *kptn^Δ5/ Δ14^* embryos featuring peaks to represent the number of tail twitches. The Y axis represents the amplitude of the tail twitches, and the X axis represents the duration of recording in seconds. (B and C) Quantification of STC behaviour. (B) Analysis of frequency in *kptn^+/+^* (n = 64 embryos), *kptn^Δ5/ Δ14^* (n = 101 embryos) and *kptn^Δ5/ Δ14^ + zkptn^WT^* (n = 69 embryos); ****p<0.0001; ns=non-significant, p>0.9999. (C) Analysis of maximum amplitude in *kptn^+/+^* (n = 64 embryos), *kptn^Δ5/ Δ14^* (n = 101 embryos) and *kptn^Δ5/ Δ14^ + zkptn^WT^* (n = 69 embryos); ****p<0.0001; ns, p=0.4220. Full-length, wild-type *zkptn* could rescue both the frequency and maximum amplitude observed in the *kptn^Δ5/ Δ14^* embryos. (D) Representative plot showing trajectories of distance travelled by *kptn^+/+^* and *kptn^Δ5/ Δ14^* embryos. (E and F) Quantification of TEER behaviour. E) Analysis of average velocity in *kptn^+/+^* (n = 23 embryos), *kptn^Δ5/ Δ14^* (n = 24 embryos) and *kptn^Δ5/ Δ14^ + zkptn^WT^* (n = 23); *p<0.05; ns, p=0.9897. (F) Analysis of maximum acceleration in *kptn^+/+^* (n = 23 embryos), *kptn^Δ5/ Δ14^* (n = 24 embryos) and *kptn^Δ5/ Δ14^ + zkptn^WT^* (n = 23); ****p<0.0001; ns, p>0.9999. Data are pooled from at least three independent biological replicates. Individual points represent single embryo; the mean is indicated by a dotted line, with error bars denoting ± SEM. Statistical significance was assessed using the Kruskal–Wallis test with Dunn’s multiple-comparison correction (B,F) or the Brown-Forsythe and Welch ANOVA test for experiments with more than two groups (C,E).

All *kptn^Δ5/ Δ14^* and *kptn^+/+^* larvae evaluated for TEER at 60 hpf elicited an escape response, suggesting an intact sensory component. However, the motor component of the escape response revealed deficits in the *kptn* null animals. In contrast to *kptn^+/+^* (6.77 ± 0.22 cm/s), the *kptn^Δ5/ Δ14^* (5.91 ± 0.20cm/s) zebrafish had reduced swimming velocity. The maximum acceleration was also decreased in *kptn^Δ5/ Δ14^* (5.71 ± 0.15m/s^2^) with respect to the *kptn^+/+^* (6.96 ± 0.20 m/s^2^), though the total distance travelled by wild-type (*kptn^+/+^*) and *kptn* null animals (*kptn^Δ5/ Δ14^*) was comparable (Figure 1D-F).

Rescue experiments were conducted by injecting mRNA of full-length zebrafish *kptn* tagged with mNeonGreen (*zkptn^WT^*) into *kptn^Δ5/ Δ14^* embryos at the one-cell stage. Embryos positive for mNeonGreen fluorescence were used to assess behaviours. Ectopic expression of *kptn* in *kptn^Δ5/ Δ14^* animals rescued both the frequency (4.99 ± 0.12 min^-1^) and intensity (10.66 ± 0.18 a.u.) of tail twitches to wild-type levels (Figure 1A-C). Similarly, in TEER, the swimming velocity (6.87 ± 0.26 cm/s) and the maximum acceleration (7.42 ± 0.31 m/s^2^) of *kptn^Δ5/ Δ14^* fish were restored to wild-type levels by full-length *zkptn^WT^* mRNA injection (Figure 1D-F). These experiments confirm that the observed behavioural deficits were due to the deletion of *kptn*.

### 2. *kptn* regulates interstitial axon branching and neuromuscular junction density

To understand the aetiology of impaired motor behaviours observed in *kptn* null animals, we assessed the development of the primary motor neurons innervating the trunk muscles. To label the motor neurons, *kptn^Δ14/ Δ14^* was crossed to Tg(mnx1:GFP), a transgenic line that expresses GFP in zebrafish motor neurons. For all the experiments the *kptn^Δ14/ Δ14^*; Tg(mnx1:GFP) was crossed with *kptn^Δ5/ Δ5^* to attain the heteroallelic condition i.e. *kptn^Δ5/ Δ14^*; Tg(mnx1:GFP).

The first primary motor neuron to extend its axon from the spinal cord and innervate the ventral musculature is the caudal primary motor neuron (CaP). The other two primary motor neurons – the middle primary (MiP) and the rostral primary (RoP)-initially follow the path taken by the CaP motor neuron (Myers et al., 1986). The *kptn* mutants, like their wild-type siblings, had somata for all three primary motor neurons in all the hemisegments of the spinal cord. These cell bodies produced axonal outgrowths that innervated the muscles, with all three primary motor neurons reaching their choice points at the same stage as their wild-type siblings, suggesting normal motor neuron outgrowth.

Motor neurons develop collateral branches from the main axonal shaft, and the development of interstitial branching can be easily observed in CaP neurons. At 26 hpf, CaP axons display multiple short filopodia-like protrusions, while by 60 hpf, a proportion of these mature into stable arbours (Myers et al., 1986). Comparing the Kptn-deficient and wild-type animals, we observed that *kptn^Δ5/ Δ14^* zebrafish (0.48 ± 0.02 protrusions/µm) displayed increased protrusion density (the number of protrusions normalised to the length of the CaP motor neuron) compared to the *kptn^+/+^* animals (0.21 ± 0.01 protrusions/µm) (Figure 2A-D). Consistent with these early protrusions representing the initiation of collateral branches, a substantially higher branch density (number of collateral branches normalised to the length of the CaP motor neuron) was observed at 60 hpf in *kptn^Δ5/ Δ14^* (0.11 ± 0.002 branches/µm) with respect to the *kptn^+/+^* (0.07 ± 0.002 branches/µm) (Figure 2E-H).

**Figure 2.**
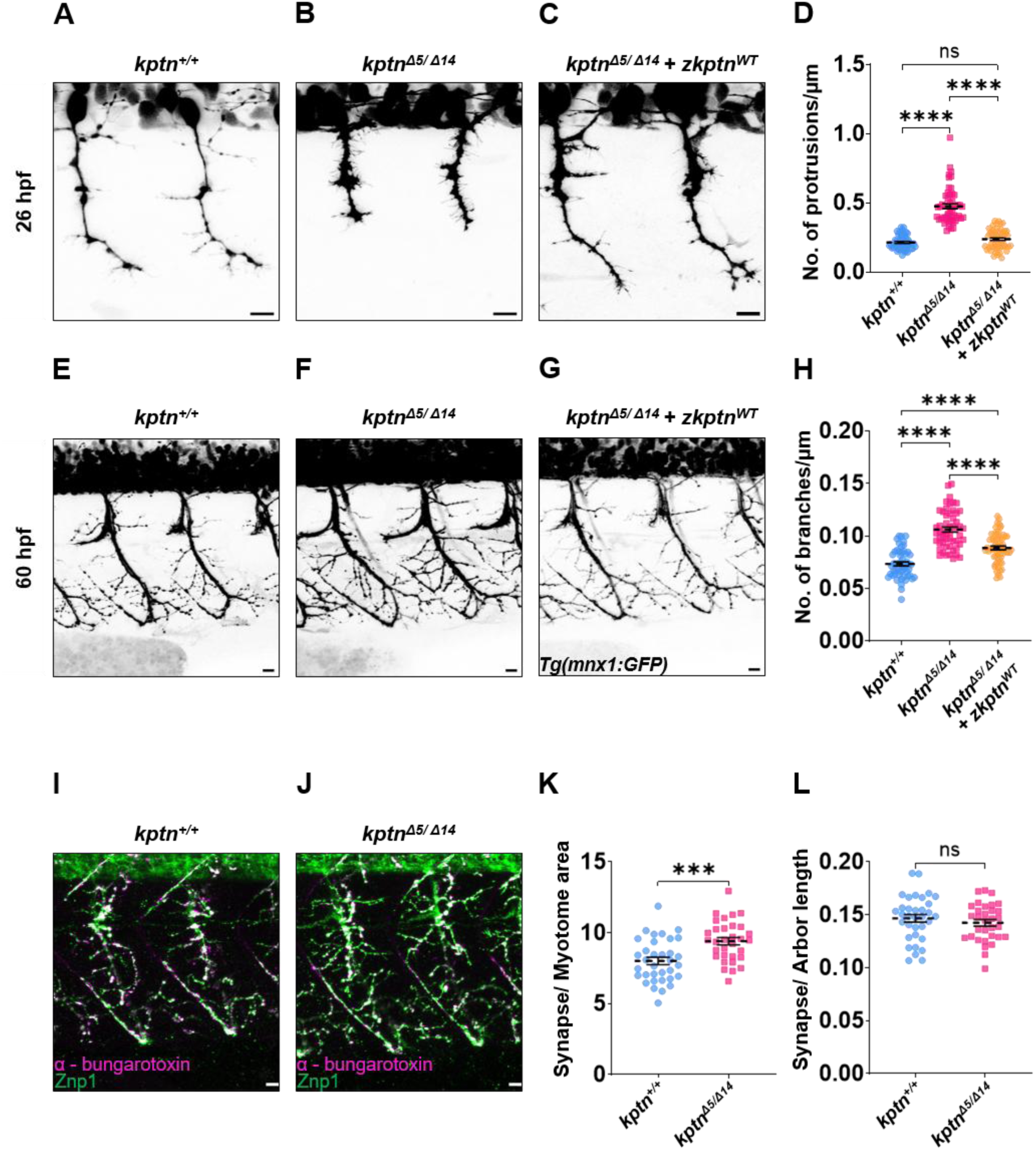
Kptn regulates motor neuron branching and NMJ density in zebrafish. (A-C) Representative micrographs of 26 hpf primary motor neurons in Tg(mnx1: GFP) zebrafish larvae of (A) *kptn^+/+^* (B) *kptn^Δ5/ Δ14^* and (C) *kptn^Δ5/ Δ14^ + zkptn^WT^.* Full-length *zkptn* mRNA was injected at the single-cell stage in *kptn^Δ5/ Δ14^* embryos. (D) Quantification of protrusion density (no. of protrusions per axon) in *kptn^+/+^* (n = 18 embryos/54 neurons), *kptn^Δ5/ Δ14^* (n = 18 embryos/54 neurons) and *kptn^Δ5/ Δ14^ + zkptn^WT^* (n = 18 embryos/54 neurons); ****p<0.0001; ns, p=0.7049. (E-G) Representative micrographs of 60 hpf primary motor neurons in Tg(mnx1: GFP) zebrafish larvae of (A) *kptn^+/+^,* (B) *kptn^Δ5/ Δ14^* and (C) *kptn^Δ5/ Δ14^ + zkptn^WT^.* Full-length *zkptn* mRNA was injected at the single-cell stage in *kptn^Δ5/ Δ14^* embryos. (H) Quantification of branch density (no. of branches per axon) in *kptn^+/+^* (n = 18 embryos/53 neurons), *kptn^Δ5/ Δ14^* (n = 18 embryos/54 neurons) and *kptn^Δ5/ Δ14^ + zkptn^WT^*(n = 21 embryos/61 neurons); ****p<0.0001. (I and J) Representative micrograph of 60 hpf embryos immunostained with a presynaptic marker znp1 and postsynaptic marker α-bungarotoxin, (I) *kptn^+/+^* (J) *kptn^Δ5/ Δ14^.* (K and L) Quantification of colocalization between znp1 and α-bungarotoxin using the SynapCountJ plugin. (K) Analysis of number of synapses per unit myotome area in *kptn^+/+^* (n = 34 neurons) and *kptn^Δ5/ Δ14^* (n = 32 neurons); ***p=0.0003. (L) Analysis of the number of synapses per unit arbour length in *kptn^+/+^* (n = 34 neurons) and *kptn^Δ5/ Δ14^* (n = 32 neurons). ns, p=0.3794. Data are pooled from at least three independent biological replicates. Individual points represent a single neuron; the mean is indicated by a dotted line, with error bars denoting ± SEM. Statistical significance was assessed using the Kruskal–Wallis test with Dunn’s multiple-comparison correction (D) and the Brown-Forsythe and Welch ANOVA test for experiments with more than two groups (H) or the Welch’s t-test for two-group comparisons (K,L). Scale bar: 10 μm (A–C, E-G, I,J).

Injection of full-length *zkptn*^WT^ mRNA in *kptn* null zebrafish partially rescued both the protrusion density at 26 hpf (0.24 ± 0.01 protrusions/µm) and the branch density at 60 hpf (0.09 ± 0.002 branches/µm).

These results reveal a novel role of Kptn in the regulation of axon collateral branching in zebrafish CaP motor neurons.

Motor neurons form synaptic contacts with the myotomes, and functional neuromuscular junctions (NMJs) develop on the zebrafish ventral musculature by 48 hpf (Egashira et al., 2018). To evaluate the NMJs, double immunostaining with presynaptic (Znp1) and postsynaptic (α-bungarotoxin) was undertaken. Colocalization of pre- and postsynaptic markers was used to score functional synapses. We observed a higher number of synapses per myotome area for *kptn^Δ5/ Δ14^* (9.41 ± 0.25) when compared to *kptn^+/+^* (8.02 ± 0.26). However, the number of synapses per unit arbour length was unchanged between *kptn^Δ5/ Δ14^* (0.14 ± 0.003) and the *kptn^+/+^* (0.15 ± 0.004) (Figure 2I-L). These data suggest that the observed increase in the number of NMJs is primarily driven by excessive branching, resulting in more elaborate arbours.

We examined muscle architecture using phalloidin staining at 60 hpf to determine whether, in addition to an excess of NMJs, structural changes in the muscles contributed to the motor deficits observed in *kptn* mutants. However, the gross muscle morphology and the organisation of F-actin bundles within the myotome for *kptn* mutants were comparable to the wild type fish (Figure S5).

Taken together, Kptn is identified as a central inhibitor of collateral branching in zebrafish motor neurons. Excessive branching in *kptn* null animals increases the NMJ density on myotomes and likely results in deregulated muscle function and motor deficits.

### 3. Kptn regulates axonal collateral branching in primary neuronal cultures

To uncover the mechanistic basis of Kptn-mediated regulation of collateral branching, we evaluated Kptn function in primary cultures of embryonic chick spinal neurons (Kundu et al., 2022, 2021; Sahasrabudhe et al., 2016). A specific translation blocking morpholino against chicken Kptn (Kptn MO) was used to knockdown endogenous Kptn protein levels (Figure S6). Morpholinos were co-transfected with a plasmid expressing GFP, and only neurons expressing GFP were selected for analysis. At 36 hours (h) *in vitro*, a marked increase in the frequency of F-actin–positive nascent axonal filopodia-like protrusions was observed in neurons transfected with Kptn MO (0.44 ± 0.02 protrusions/µm) as compared to those transfected with a non-specific control morpholino (Ctrl MO; 0.28 ± 0.02 protrusion/µm) (Figure 3A-D). This increase in protrusion density was rescued (0.29 ± 0.02 protrusions/µm) by co-transfection of Kptn MO with a morpholino-resistant, full-length wild-type Kptn construct (Kptn^WT^-GFP), underscoring the specificity of the Kptn MO (Figure 3A-D). Interestingly, there was no difference in growth cone filopodia numbers between the Ctrl MO-treated and Kptn MO-treated neurons (Figure S7). These data suggest that Kptn specifically restricts the formation of axonal filopodia-like protrusions that initiate the development of axonal collateral branches.

**Figure 3.**
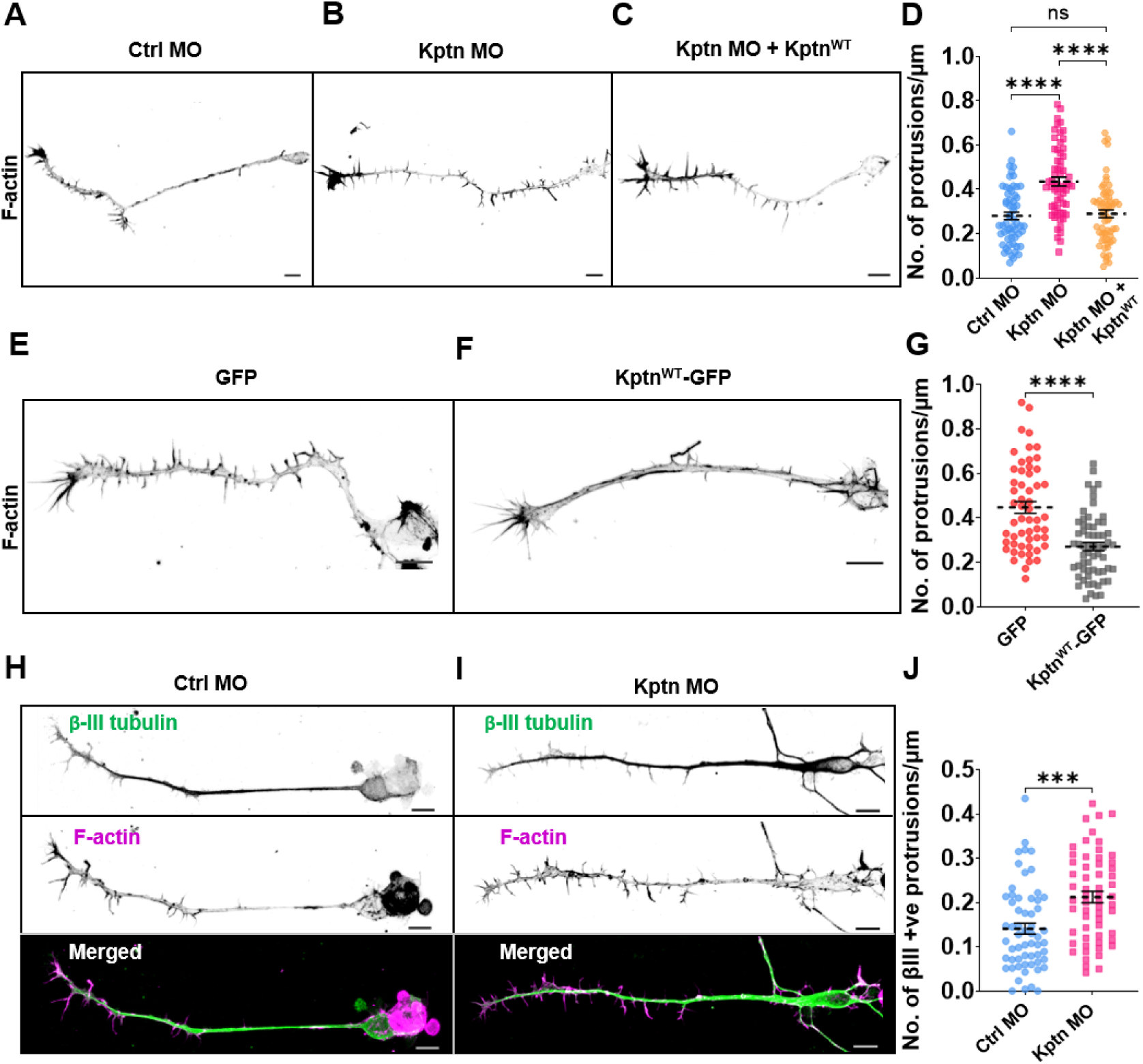
Kptn restricts axonal collateral branching in cultured neurons. (A–C) Representative images of cultured neurons transfected with Ctrl MO + pCAG-GFP (A), Kptn MO + pCAG-GFP (B), or Kptn MO + pCAG-Kptn^WT^-GFP (C). Neurons were fixed at 36 h post-plating and stained with Alexa Fluor 568–conjugated phalloidin. (D) Quantification of axonal protrusion density at 36 h post-plating (Ctrl MO, n = 59 axons; Kptn MO, n = 62 axons; Kptn MO + C, n = 62 axons; ****p<0.0001). (E, F) Representative phalloidin-stained neurons overexpressing pCAG-GFP (E) or pCAG- Kptn^WT^-GFP (F) at 36 h post-plating. (G) Quantification of axonal protrusion density following pCAG-GFP (n = 55 axons) or pCAG- Kptn^WT^-GFP (n = 62 axons) overexpression (****p<0.0001). (H, I) Representative neurons treated with Ctrl MO (H) or Kptn MO (I) together with pCAG-GFP, fixed at 72 h post-plating and stained for βIII-tubulin and F-actin. (J) Quantification of βIII-tubulin–innervated axonal protrusion density at 72 h post-plating (Ctrl MO, n = 59 axons; Kptn MO, n = 59 axons; ***p=0.0001). Data are pooled from at least three independent biological replicates. Individual points represent single axons; the mean is indicated by a dotted line, with error bars denoting ± SEM. Statistical significance was assessed using the Kruskal–Wallis test with Dunn’s multiple-comparison correction for experiments with more than two groups (D) or the Mann–Whitney *U* test for two-group comparisons (G, J). Scale bar: 10 μm (A–C, E, F, H, I).

Next, we overexpressed GFP-tagged Kptn (Kptn^WT^-GFP) in cultured neurons. A substantial decrease in protrusion density was observed in Kptn overexpressing neurons (0.27 ± 0.02 protrusions/µm) as compared to control neurons transfected with the empty backbone plasmid (0.45 ± 0.03 protrusions/µm) (Figure 3E-G). This gain-of-function experiment shows that Kptn is sufficient for inhibiting the formation of nascent axonal protrusions.

Entry and capture of microtubules in axonal protrusions is necessary for the stabilisation of the filopodia-like processes and their subsequent transition to a stable collateral branch (Dent and Kalil, 2001; Ketschek et al., 2015; Kundu et al., 2022).

Thus, to evaluate if the increase in axonal protrusions due to Kptn knockdown translates to mature axon interstitial branches, we scored the density of F-actin rich axonal projections that were also positive for βIII-tubulin at 72 h *in vitro*. Kptn MO-transfected neurons exhibited a significantly higher density of ꞵIII-tubulin-positive stable protrusions (0.21 ± 0.01 protrusions/µm) relative to the Ctrl MO-treated neurons (0.14 ± 0.01 protrusions/µm) (Figure 3H-J).

Collectively, these results indicate that Kptn activity is necessary and sufficient to restrict the initiation of axonal protrusions and, in turn, regulate axonal collateral branching. This function is conserved in zebrafish motor neurons and chick spinal neurons.

### 4. Kptn regulates collateral protrusion initiation by restricting the maturation axonal F-actin patches

Axonal actin patches are localised, transient assemblies of branched F-actin that form along the axon shaft and serve as precursor structures for axonal protrusions and subsequent collateral branches. Following their formation, actin patches exist in a dynamic continuum of assembly and disassembly cycles regulated by the activities of actin-binding proteins that either promote or restrict actin filament growth (Armijo-Weingart and Gallo, 2017; Gallo, 2011; Kundu et al., 2022). Progression towards mature, stabilised states, as reflected in increased patch size and lifetime, favours the transition of these F-actin patches into protrusions. Conversely, patch dissipation precludes the initiation of axonal protrusions (Gallo, 2024; Kundu et al., 2022). Recent findings from (Dutta et al., 2026) demonstrate that Kptn associates with F-actin barbed ends in *in vitro* reconstitution assays and exhibits filament capping activity by restricting filament elongation. We therefore investigated the premise that Kptn regulates axonal F-actin patch dynamics to influence collateral branching.

To test if Kptn is associated with the F-actin patches that give rise to axonal protrusions, neurons were co-transfected with Kptn^WT^-GFP and the F-actin probe, F-tractin–mCherry, and imaged using time-lapse spinning-disk confocal microscopy. Typically, the appearance of an axonal F-actin patch coincided with a localised enrichment of Kptn^WT^-GFP at the F-actin patch. The Kptn enrichment concurrently increased with the increase in the F-actin patch size, and if an actin-rich deformation was generated, the Kptn signal extended into the emerging protrusion along with F-actin (Figure 4A). These data suggest that Kptn is recruited to F-actin patches generating axonal protrusions.

**Figure 4.**
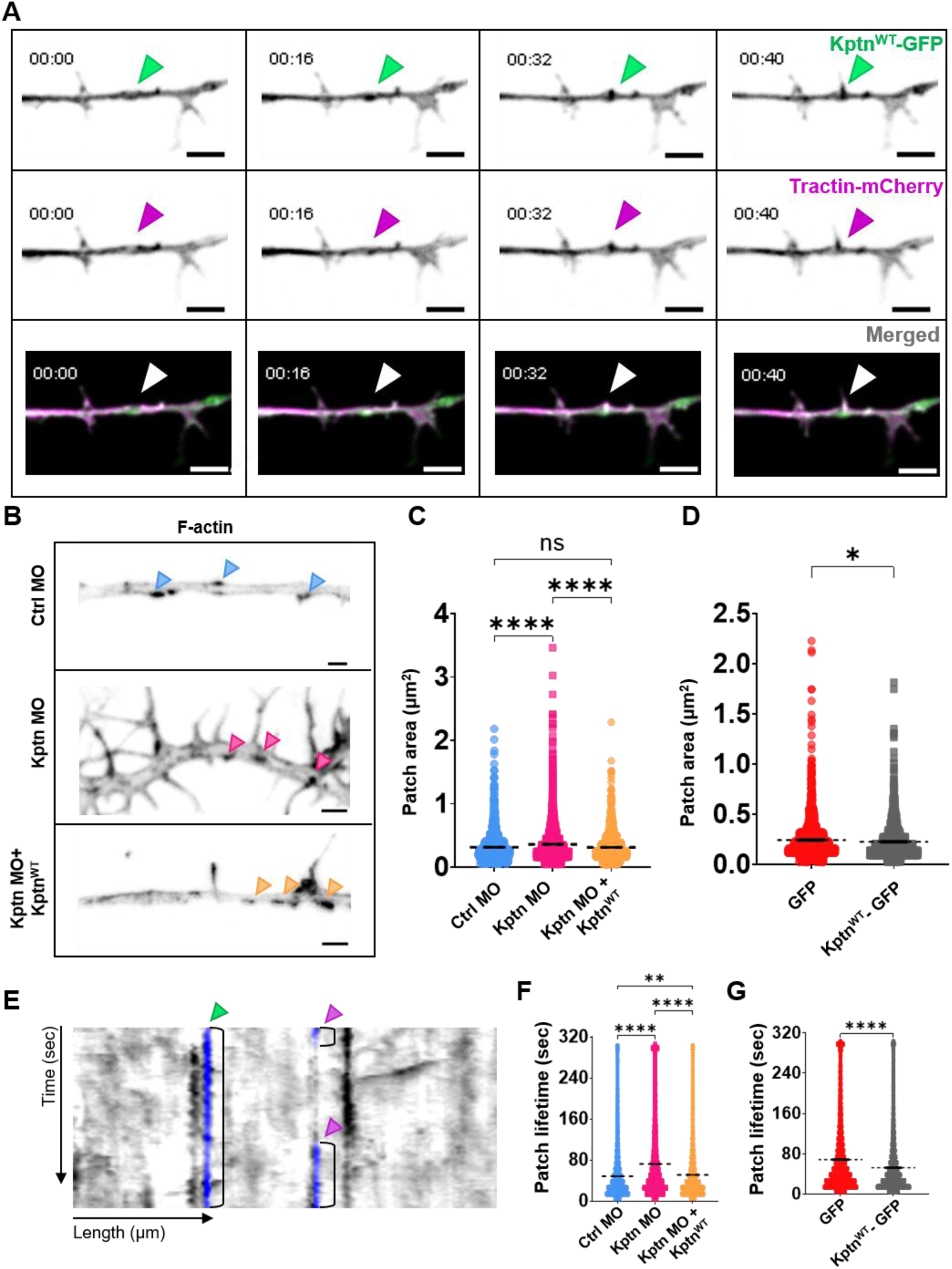
Kptn controls axonal protrusion initiation by limiting F-actin patch maturation. (A) Montage from time-lapse image sequence of a neuron co-transfected with pCAG-F-tractin-mCherry (Tractin-mCherry) and pCAG-Kptn^WT^-GFP (Kptn^WT^-GFP). The top row shows the Kptn^WT^ channel, the middle row shows the F-tractin channel, and the bottom row shows the overlay of both channels. Kptn (green) and F-actin (magenta) signals showed little spatial overlap along the axonal shaft at time 00:00. However, upon the appearance of a discrete actin patch at 00:16 s, a localised enrichment of Kptn^WT^-GFP was observed at the same site (marked by an arrowhead). As the time progressed, both the actin patch and the associated Kptn enrichment increased concurrently in size and fluorescence intensity (00:32 s). Subsequently, the actin patch gave rise to an axonal protrusion, with the Kptn signal extending into the emerging protrusion. (B) Representative images of axons stained with Alexa Fluor 568–conjugated phalloidin and transfected with Ctrl MO + pCAG-GFP, Kptn MO + pCAG-GFP, or Kptn MO + pCAG-Kptn^WT^-GFP. Arrowheads mark axonal F-actin patches under the indicated conditions. (C, D) Quantification of axonal actin patch area. (C) Patch area corresponding to the conditions shown in (B): Ctrl MO + pCAG-GFP (n = 1,206 patches/58 axons), Kptn MO + pCAG-GFP (n = 2,356 patches/62 axons), and Kptn MO + pCAG-Kptn^WT^-GFP (n = 1,438 patches/62 axons); ****p<0.0001. (D) Patch area following overexpression of pCAG-GFP (n = 1,566 patches/55 axons) or pCAG-Kptn^WT^-GFP (n = 1,360 patches/63 axons); *p=0.0367. (E) Kymograph generated from a representative time-lapse sequence of a neuron transfected with pCAG-F-tractin-mCherry to assess actin patch dynamics in live. Representative actin patches are marked by vertical blue lines, the lengths of which represent patch lifetimes. The F-actin patch labelled by magenta arrowheads undergoes repeated appearance and disappearance at the same axonal location (“blinking”), whereas the F-actin patch labelled by green arrowhead remains stable over the imaging period. (F, G) Quantification of actin patch lifetime under the indicated conditions, measured from F-tractin–mCherry kymographs: (F) Ctrl MO + pCAG-GFP (n = 6,923 patches/57 axons), Kptn MO + pCAG-GFP (n = 8,463 patches/67 axons), and Kptn MO + pCAG-Kptn^WT^-GFP (n = 8,476 patches/56 axons); (G) pCAG-GFP (n = 8,323 patches/55 axons) and pCAG-Kptn^WT^-GFP (n = 8,174 patches/59 axons); ****p<0.0001, **p=0.0013. Data are pooled from at least three independent biological replicates. Individual points represent single actin patches; the mean is indicated by a dotted line, with error bars denoting ± SEM. Statistical significance was assessed using the Kruskal–Wallis test with Dunn’s multiple-comparison correction for experiments with more than two groups (C, F) or the Mann–Whitney *U* test for two-group comparisons (D, G). Scale bars: 3 μm in (A), and 2 μm in (B).

To analyse F-actin patch stability, we modulated Kptn levels by either knocking it down using Kptn MO or by overexpressing Kptn^WT^ and evaluated F-actin patches using fluorophore-conjugated Phalloidin in fixed neurons. Kptn depletion resulted in a considerable increase in mean patch area (0.36 ± 0.01 µm^2^) as compared to the controls (0.31 ± 0.01 µm^2^) (Figure 4B and C). Conversely, overexpression of Kptn reduced the patch area (0.22 ± 0.01 µm^2^) in comparison to control neurons (0.24 ± 0.01 µm^2^) (Figure 4D). Rescue experiments expressing full-length Kptn^WT^ in Kptn-depleted neurons restored the mean patch area (0.31 ± 0.01 µm^2^) to control levels (Figure 4B and C). Together, these results suggest that Kptn negatively regulates F-actin-patch size.

We next examined whether perturbation of Kptn also affected F-actin patch lifetime. Cultured neurons expressing the F-actin probe F-tractin-mCherry were subjected to Kptn knockdown or overexpression and imaged live. Kymograph analysis of time-lapse sequences was used to quantify the lifetime of individual F-tractin–positive actin patches across different conditions (Figure 4E). Knockdown of Kptn significantly increased the mean lifetime of axonal F-actin patches (72.30 ± 0.80 seconds) compared to control neurons (48.51 ± 0.60 seconds) (Figure 4F). Expression of full-length Kptn^WT^ in Kptn knockdown neurons restored patch lifetime to control levels (51.12 ± 0.57 seconds) (Figure 4F). Conversely, overexpression of Kptn reduced actin patch lifetime: from 68.27 ± 0.77 seconds in control neurons to 52.18 ± 0.63 seconds in Kptn-overexpressing neurons (Figure 4G).

Taken together, and consistent with its proposed role in restricting F-actin elongation by barbed end capping (Dutta et al., 2026), these experiments implicate Kptn in constraining the growth and maturation of axonal F-actin patches. As a consequence, Kptn regulates axon interstitial branching and axonal arbourisation.

### 5. F-actin barbed-end binding activity of Kptn dictates actin patch dynamics and collateral branching

A conserved positively charged amino acid residue (R59) in Kptn has been shown to be essential for F-actin-binding and restriction of barbed-end elongation (Dutta et al., 2026) (Figure S1). We substituted the arginine in the 59th position of Kptn to aspartate (Kptn^R59D^) to directly test if restriction of actin filament growth regulated interstitial branch initiation and F-actin patch dynamics. As shown previously (Figure 3A-D; 4B,C and F), expression of Kptn^WT^ in Kptn MO-transfected neurons rescued both the protrusion and actin patch phenotypes in cultured neurons. In contrast, expression of the Kptn^R59D^ mutant in Kptn knockdown neurons failed to restore protrusion density, which remained statistically comparable to that observed in Kptn MO–treated neurons alone (0.50 ± 0.02 protrusions/µm) (Figure 5 A and C). Similarly, Kptn^R59D^ failed to rescue F-actin patch lifetime (69.05 ± 0.72 seconds) or F-actin patch area (0.37 ± 0.01s µm^2^) (Figure 5 B,D-F).

**Figure 5.**
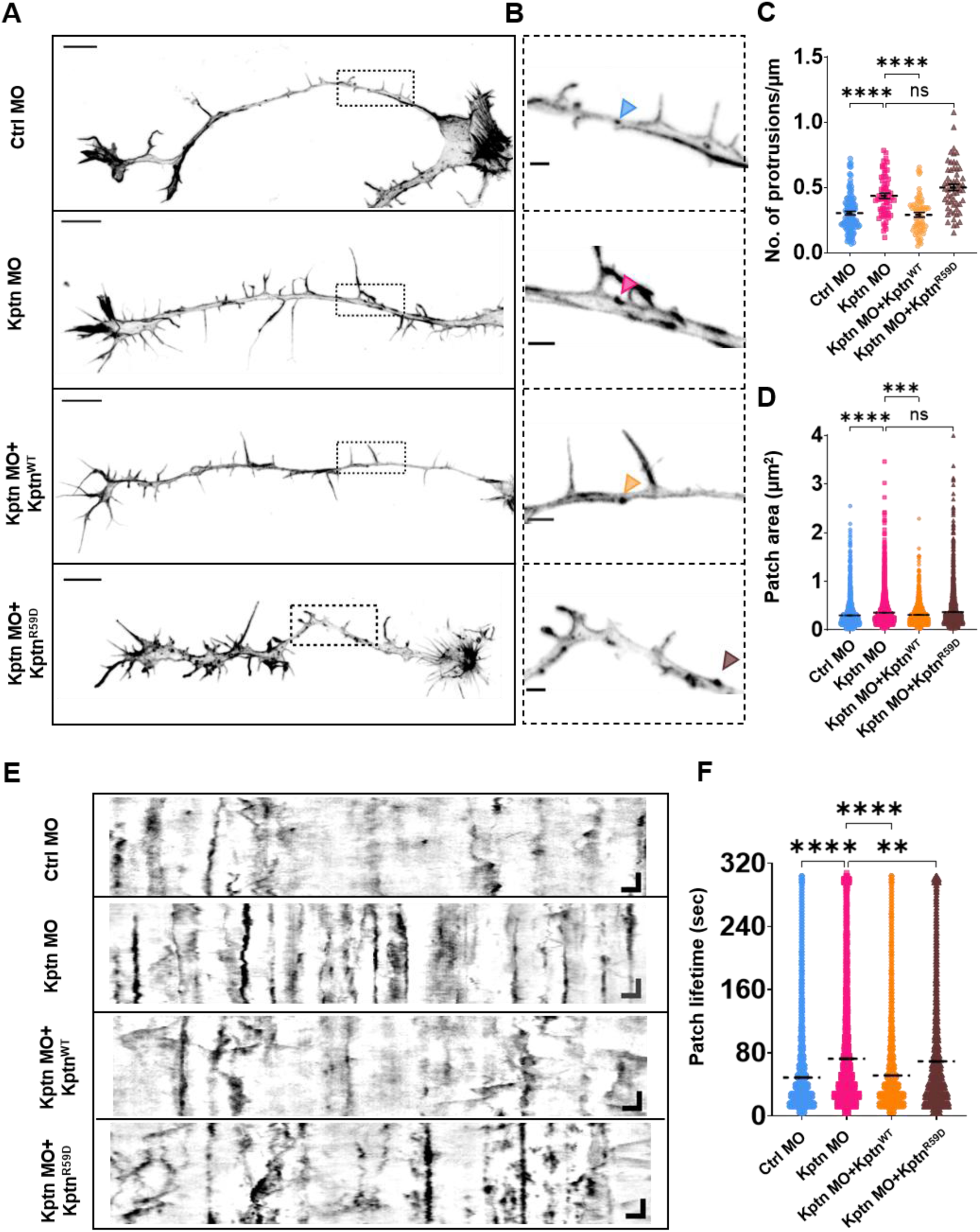
Kptn regulates axonal patch stability and protrusion density through F-actin-binding and inhibition of barbed-end elongation. (A) Representative cultured neurons transfected with the indicated constructs and stained with Phalloidin to visualize F-actin. (B) Enlarged views of boxed regions in (A), highlighting axonal actin patches (arrowheads). (C) Axonal protrusion density under the indicated conditions: Ctrl MO + pCAG-GFP (n = 108 axons), Kptn MO + pCAG-GFP (n = 62 axons), Kptn MO + pCAG-Kptn^WT^-GFP (n = 62 axons), and Kptn MO + pCAG-Kptn^R59D^-GFP (n = 57 axons); ****p<0.0001. (D) Corresponding analysis of actin patch area: Ctrl MO + pCAG-GFP (n = 2,559 patches/108 axons), Kptn MO + pCAG-GFP (n = 2,356 patches/62 axons), Kptn MO + pCAG- Kptn^WT^-GFP (n = 1,438 patches/62 axons), and Kptn MO + pCAG-Kptn^R59D^-GFP (n = 2,934 patches/57 axons); ****p<0.0001, ***p=0.0001. (E) Representative F-tractin–mCherry kymographs depicting actin patch dynamics under the indicated transfection conditions. (F) Summary of actin patch lifetimes quantified from F-tractin–mCherry kymographs: Ctrl MO + pCAG-GFP (n = 6,923 patches/57 axons), Kptn MO + pCAG-GFP (n = 8,463 patches/67 axons), Kptn MO + pCAG-Kptn^WT^-GFP (n = 8,476 patches/56 axons), and Kptn MO + pCAG-Kptn^R59D^-GFP (n = 9,730 patches/65 axons); ****p<0.0001, **p=0.0036. Data from at least three independent biological replicates were combined for analysis. Each data point corresponds to a single neuron in (C) or an individual actin patch in (D, F). Mean is indicated by a dotted line, with error bars representing ± SEM. Statistical comparisons were performed using the Kruskal–Wallis test followed by Dunn’s multiple-comparison correction. Scale bars: 10 μm (A), 2 μm (B), and 2 μm (horizontal) X 60 s (vertical) in (E).

In zebrafish Kptn, the residue equivalent to mammalian R59 is lysine at position 55 (K55) (Figure S1). We therefore substituted at the 55^th^ amino acid position with aspartic acid of zebrafish Kptn (*zkptn^K55D^*). Consistent with the results obtained *in vitro* (Figure 5), the protrusion density of CaP motor neurons at 26 hpf for the *zkptn^K55D^* injected *kptn^Δ5/ Δ14^* embryos (0.40 ± 0.01 protrusions/µm) continued to remain higher compared to the *kptn^+/+^* (0.21 ± 0.01 protrusions/µm) (Figure 6A-D). Similarly, at 60 hpf, the increased branch density in *kptn^Δ5/ Δ14^* could not be rescued by injecting *zkptn^K55D^* (0.12 ± 0.002 branches/µm) (Figure 6E-H).

**Figure 6.**
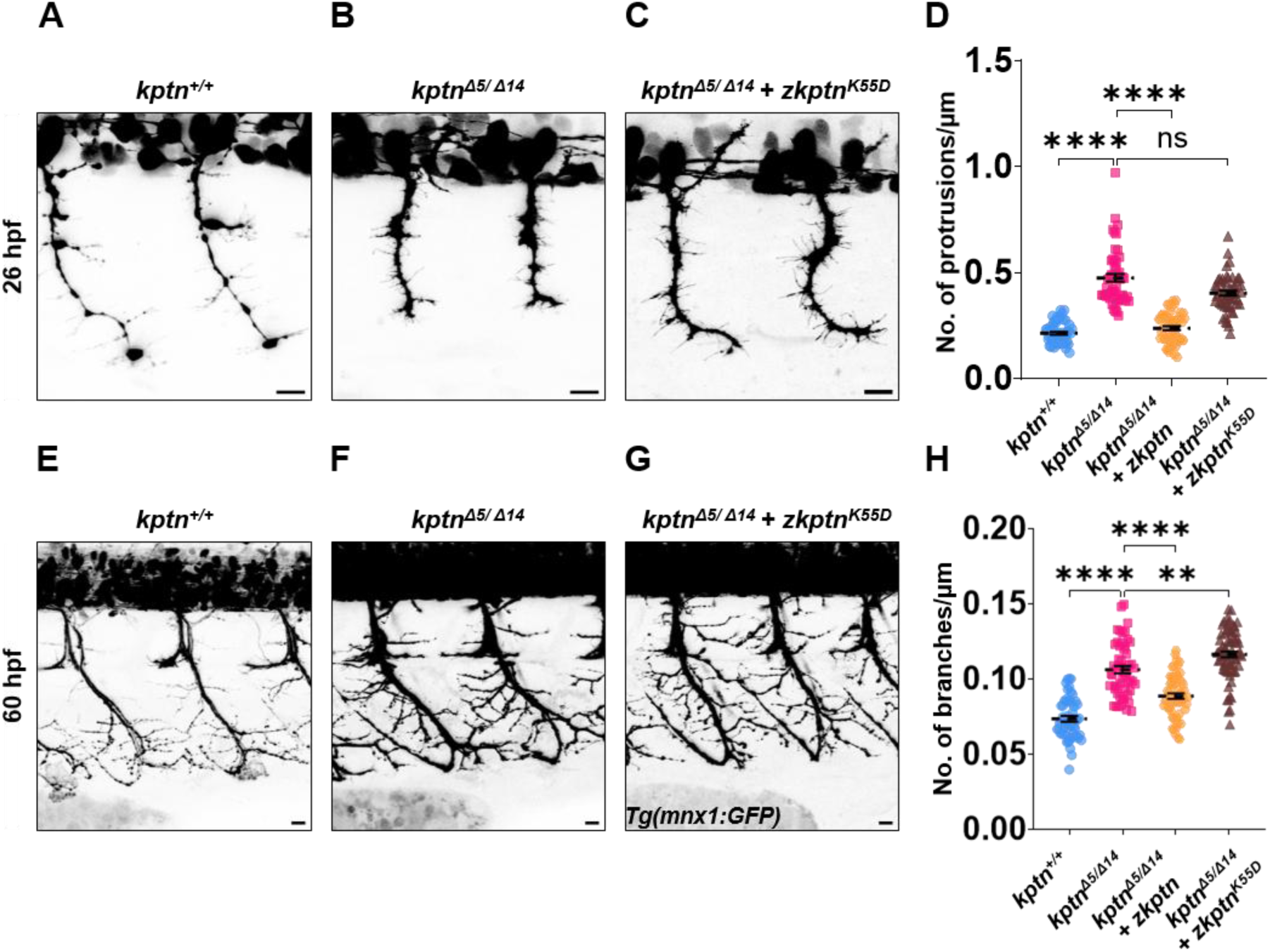
Kptn regulates zebrafish motor neuron branching through its F-actin barbed end binding. (A-C) Representative micrographs of 26 hpf primary motor neurons in Tg(mnx1: GFP) zebrafish larvae of (A) *kptn^+/+^* (B) *kptn^Δ5/ Δ14^* and (C) *kptn^Δ5/ Δ14^ + zkptn^K55D^. zkptn^K55D^* mRNA was injected at the single cell stage in *kptn^Δ5/ Δ14^* embryos. (D) Quantification of protrusion density (no. of protrusions per axon) in *kptn^+/+^* (n = 18 embryos/54 neurons), *kptn^Δ5/ Δ14^* (n = 18 embryos/54 neurons) and *kptn^Δ5/ Δ14^ + zkptn^K55D^* (n = 18 embryos/54 neurons); ****p<0.0001; ns, p=0.3827. (E-G) Representative micrographs of 60 hpf primary motor neurons in Tg(mnx1: GFP) zebrafish larvae of (A) *kptn^+/+^* (B) *kptn^Δ5/ Δ14^* and (C) *kptn^Δ5/ Δ14^ + zkptn^K55D^. zkptn^K55D^* mRNA was injected at the single cell stage in *kptn^Δ5/ Δ14^* embryos. (H) Quantification of branch density (no. of branches per axon) in *kptn^+/+^* (n = 18 embryos/53 neurons) and *kptn^Δ5/ Δ14^* (n = 18 embryos/54 neurons) and *kptn^Δ5/ Δ14^ + zkptn^K55D^* (n = 24 embryos/71 neurons); ****p<0.0001; **p=0.0094. Data are pooled from at least three independent biological replicates. Individual points represent single neuron; the mean is indicated by a dotted line, with error bars denoting ± SEM. Statistical significance was assessed using the Kruskal–Wallis test with Dunn’s multiple-comparison correction (D) and the Brown-Forsythe and Welch ANOVA test for experiments with more than two groups (H). Scale bar: 10 μm (A–C, E-G).

Together, these results indicate that F-actin barbed-end binding and resultant inhibition of filament elongation by Kptn are required for its ability to regulate F-actin patch and axonal protrusion phenotypes both *in vitro* and *in vivo*.

### 6. Functional antagonism between Kptn and Fmn2 determine actin-patch dynamics and, in turn, axonal collateral branching

Fmn2, a formin family F-actin nucleator/elongator, has been established as a positive regulator of axonal F-actin patches and, consequently, axonal collateral branching both in cultured neurons and in zebrafish (Gallo, 2011; Kundu et al., 2022; Nagar et al., 2021). As F-actin patch maturation appears to involve regulation of F-actin barbed end dynamics we tested if unrestricted F-actin elongation upon Kptn loss can be mitigated by limiting the opposing elongating activity of Fmn2.

To address this, we depleted both Kptn and Fmn2 in cultured neurons and quantified the protrusion phenotype. As reported previously (Kundu et al., 2022; Nagar et al., 2021), knockdown of Fmn2 alone reduced protrusion density (0.23 ± 0.02 protrusions/µm) as compared to the control conditions (0.31 ± 0.01 protrusions/µm). As demonstrated earlier, Kptn depletion led to an increase in protrusion density (0.44 ± 0.02 protrusions/µm) (Figure 3 A-D; 7C). However, concurrent depletion of both Kptn and Fmn2 was sufficient (0.31 ± 0.02 protrusions/µm) to rescue the hyperbranching phenotype observed in neurons treated with Kptn MO alone (Figure 7C). To determine whether the regulation of axonal branching by opposing activities of *kptn* and *fmn2b* (the functional Fmn2 ortholog in zebrafish) was conserved *in vivo*, we generated double homozygous mutant *fmn2b ^Δ7/ Δ7^*; *kptn^Δ14/ Δ14^* in the Tg (mnx1:GFP) transgenic background to quantify protrusion density and branch density of motor neurons at 26 hpf and 60 hpf, respectively. We observed that the double homozygous mutants *fmn2b ^Δ7/ Δ7^; kptn^Δ14/ Δ14^* have comparable protrusion density (0.30 ± 0.01 protrusions/µm) and branch density (0.09 ± 0.002 branches/µm) with respect to the wild-type animals (0.31 ± 0.01 protrusions/µm) at 26 hpf and (0.08 ± 0.002 branches/µm) at 60 hpf (Figure 7A and B).

**Figure 7.**
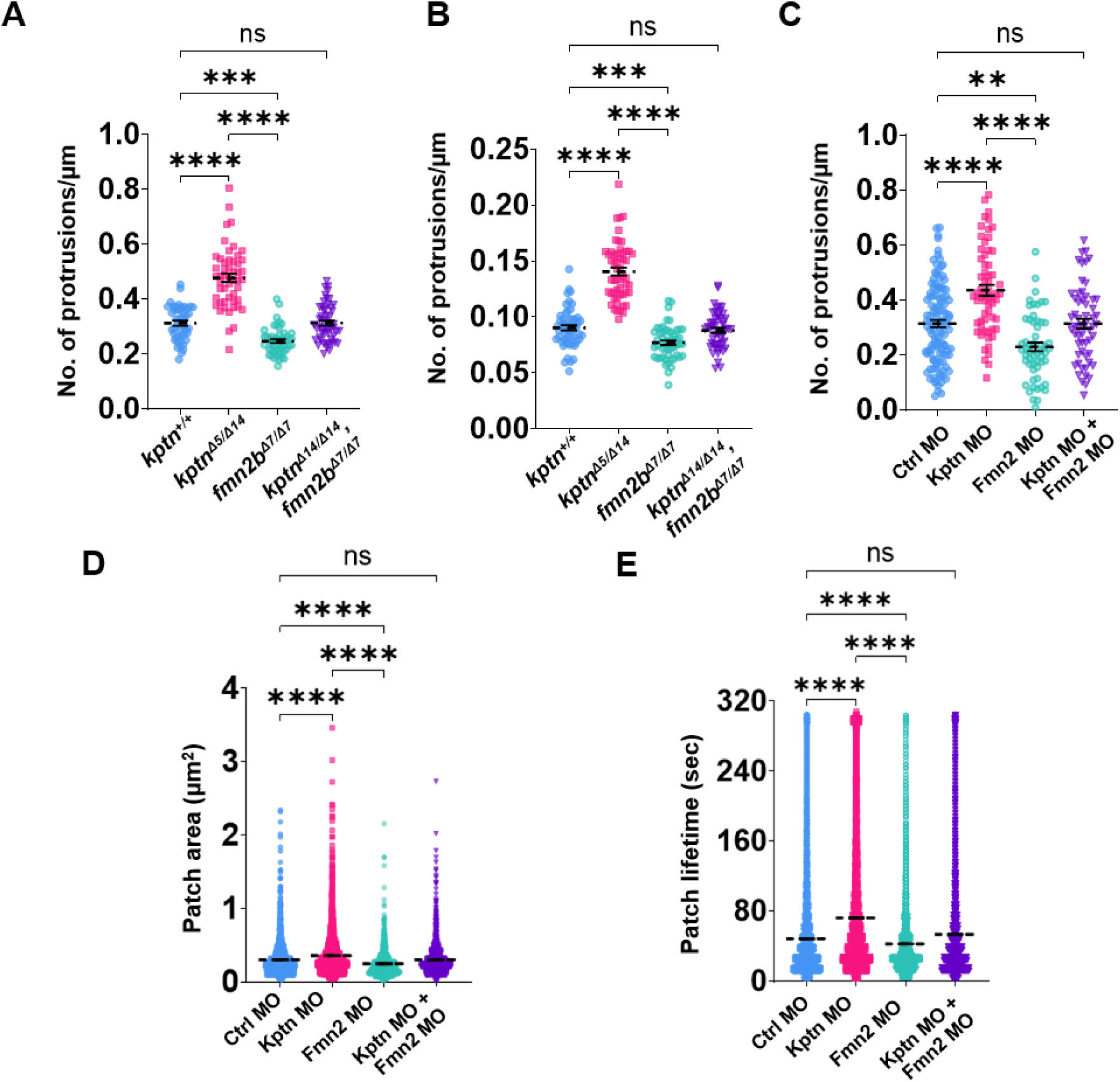
Opposing activities of Kptn and Fmn2 determine actin-patch stability and axonal collateral branching. (A) Quantification of protrusion density at 26 hpf in *kptn^+/+^* (n = 16 embryos/46 neurons), *kptn^Δ5/ Δ14^* (n = 18 embryos/51 neurons), *fmn2b ^Δ7/ Δ7^* (n = 16 embryos/48 neurons) and *fmn2b ^Δ7/ Δ7^*, *kptn^Δ14/ Δ14^* (n = 18 embryos/ 53 neurons); ****p<0.0001; ***p=0.0006; ns, p>0.9999. (B) Quantification of branch density at 60 hpf in *kptn^+/+^* (n = 18 embryos/53 neurons), *kptn^Δ5/ Δ14^* (n = 18 embryos/54 neurons), *fmn2b ^Δ7/ Δ7^* (n = 18 embryos/53 neurons) and *fmn2b ^Δ7/ Δ7^*, *kptn^Δ14/ Δ14^* (n = 18 embryos/51neurons); ****p<0.0001; ***p=0.0004; ns, p=0.9767. (C) Axonal protrusion density measured under the indicated transfection conditions: Ctrl MO + pCAG-GFP (n = 116 axons), Kptn MO + pCAG-GFP (n = 62 axons), Fmn2 MO + pCAG-GFP (n = 57 axons), and combined Kptn MO + Fmn2 MO + pCAG-GFP (n = 58 axons); ****p<0.0001, **p=0.0023. (D) Actin patch area quantified from neurons subjected to the same transfection conditions: Ctrl MO + pCAG-GFP (n = 1,754 patches/81 axons), Kptn MO + pCAG-GFP (n = 2,356 patches/62 axons), Fmn2 MO + pCAG-GFP (n = 830 patches/59 axons), and Kptn MO + Fmn2 MO + pCAG-GFP (n = 1,229 patches/59 axons); ****p<0.0001. (E) Actin patch lifetime measured under the same conditions: Ctrl MO (n = 6,923 patches/57 axons), Kptn MO (n = 8,463 patches/65 axons), Fmn2 MO (n = 10,572 patches/54 axons), and combined Kptn MO + Fmn2 MO (n = 8,977 patches/43 axons); ****p<0.0001. Data represent pooled measurements from at least three independent biological replicates. Individual points correspond to single neurons in (A-C) and single actin patches in (D, E); the mean is indicated by a dotted line, with error bars denoting ± SEM. Statistical significance was assessed using the Kruskal–Wallis test with Dunn’s multiple-comparison correction (A, C-E) and the Brown-Forsythe and Welch ANOVA test for experiments with more than two groups (B).

Furthermore, dual knockdown of Kptn and Fmn2 in cultured neurons also restored actin patch area (0.30 ± 0.006 µm^2^) and patch lifetime (53.60 ± 0.61 seconds) to control levels (0.30 ± 0.005 µm^2^ and 48.51 ± 0.60 seconds, respectively) (Figure 7 D and E).

Collectively, these results indicate that the opposing functions of Kptn and Fmn2 at the F-actin barbed end regulate axonal F-actin dynamics and, consequently, the initiation of axonal collateral branching.

### 7. Regulation of axonal branching in the zebrafish motor neurons is independent of mTORC1 signalling

Kptn has been identified as a component of the KICSTOR complex that negatively regulates mTORC1 signalling (Lupton et al., 2026; Teng et al., 2025; Wolfson et al., 2017). The literature suggests that knocking out the *kptn* gene in mice leads to mTORC1 hyperactivation (Levitin et al., 2023). mTORC1 hyperactivation drives divergent axonal branching phenotypes, manifesting as either excessive hyper-branching in some contexts or hypo-branching in others (Amegandjin et al., 2021; Carlin et al., 2018; LaSarge et al., 2015; Proietti Onori et al., 2021). It thus remains possible that hyperactivated mTORC1 in *kptn* mutants contributes to the axonal branching phenotypes observed.

To investigate whether mTORC1 is hyperactivated in the *kptn* zebrafish knockouts at stages relevant for early nervous system development, we evaluated the phosphorylation state of ribosomal S6 (p-RS6) protein, a downstream substrate of mTORC1 signalling. Immunoblots of proteins extracted from 3 dpf and 5 dpf *kptn^+/+^* and *kptn^Δ5/ Δ14^* larvae, did not reveal any differences in the levels of phosphorylated RS6 (Figure 8A-C). The KICSTOR complex belongs to the nutrient-sensing arm of the mTORC1 signalling pathway. Thus, nutrient-abundant conditions (due to the presence of larval yolk in 3 dpf and 5 dpf larvae) may not reveal mTORC1 hyperactivity in *kptn* mutants. We therefore starved 6 dpf larvae (yolk is fully absorbed by this stage) for 24 h and evaluated p-RS6 at 7 dpf. Starvation revealed no difference in mTORC1 levels in *kptn^(Δ5/Δ14)^* when compared to *kptn^+/+^* (Figure 8A and D).

**Figure 8.**
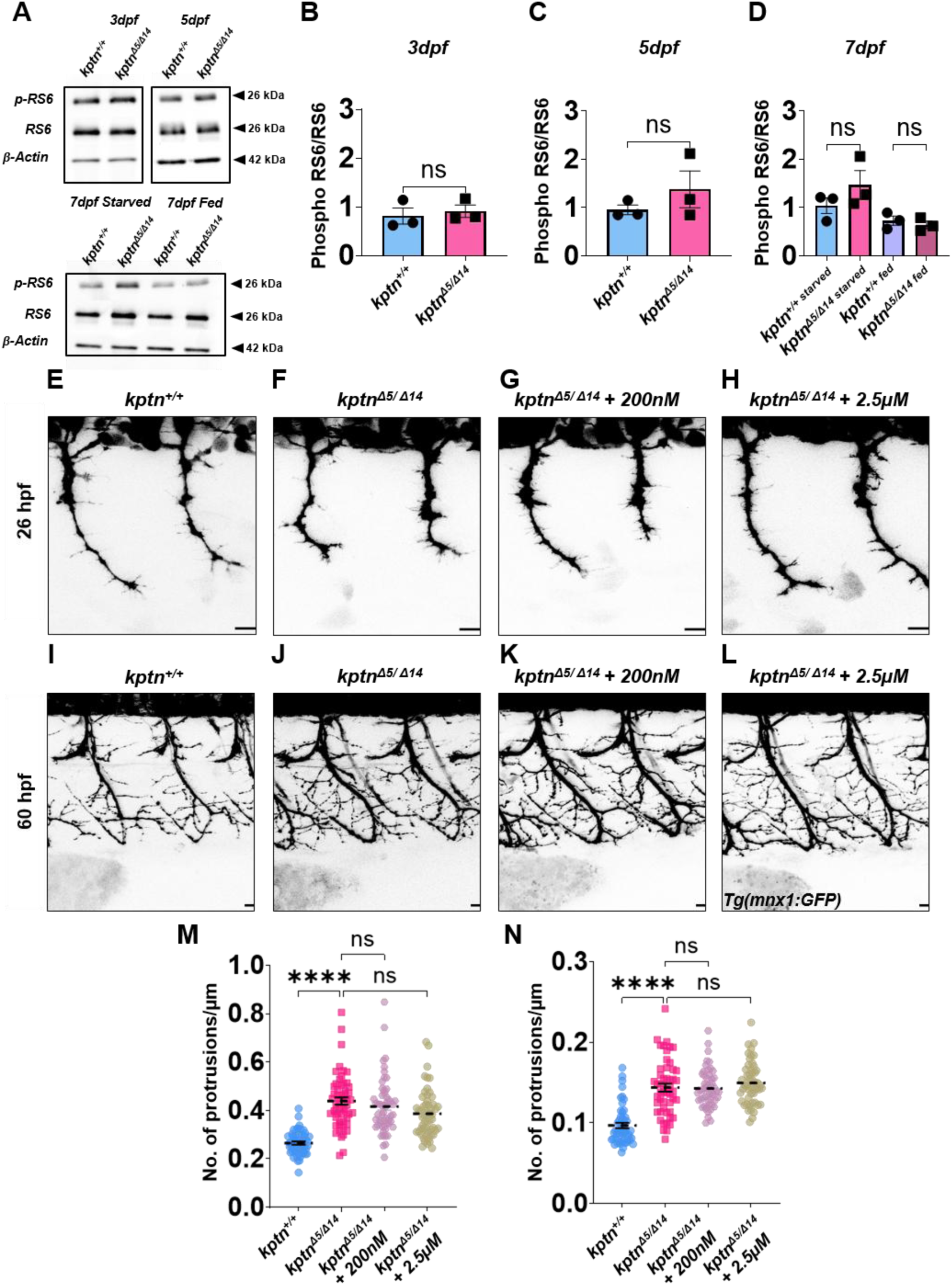
Kptn regulates motor neuron branching independent of mTORC1 signalling. (A) Representative immunoblots for p-RS6 and RS6 protein in 3 dpf, 5 dpf and 7 dpf embryos. (B) Quantification of p-RS6 level at 3 dpf showing no increase in activity between *kptn^+/+^* (n = 8 embryos/biological replicate), and *kptn^Δ5/ Δ14^* (n = 8 embryos/ biological replicate) (C) Quantification of p-RS6 level at 5 dpf showing no increase in activity between *kptn^+/+^* (n = 8 embryos/1 biological replicate) and *kptn^Δ5/ Δ14^* (n = 8 embryos/biological replicate). (D) Quantification of p-RS6 level at 7 dpf for two conditions – starved and fed - showing no increase in activity between starved *kptn^+/+^* (n = 8 embryos/1 biological replicate) and starved *kptn^Δ5/ Δ14^* (n = 8 embryos/1 biological replicate) and fed *kptn^+/+^* (n = 8 embryos/1 biological replicate) and fed *kptn^Δ5/ Δ14^* (n = 8 embryos/1 biological replicate). (E-H) Representative micrographs of 26 hpf primary motor neurons in Tg(mnx1: GFP) zebrafish larvae of (E) *kptn^+/+^* (*F*) *kptn^Δ5/ Δ14^* (G) *kptn^Δ5/ Δ14^ +* 200 nM Rapamycin and (H) *kptn^Δ5/ Δ14^ +* 2.5 µM Rapamycin. Rapamycin treatment employed two strategies – a chronic treatment with low concentration (200 nM for 16 h for 26 hpf and 32 h for 60 hpf) and an acute treatment with high concentration (2.5 µM 4 h). (I-L) Representative micrographs of 60 hpf primary motor neurons in Tg(mnx1: GFP) zebrafish larvae of of (I) *kptn^+/+^* (J) *kptn^Δ5/ Δ14^* (K) *kptn^Δ5/ Δ14^ +* 200 nM Rapamycin and (L) *kptn^Δ5/ Δ14^ +* 2.5 µM Rapamycin. (M) Analysis of protrusion density at 26 hpf in *kptn^+/+^* (n = 18 embryos/53 neurons), *kptn^Δ5/ Δ14^* (n = 18 embryos/54 neurons), *kptn^Δ5/ Δ14^* + 200 nM Rapamycin (n = 18 embryos/55 neurons) and *kptn^Δ5/ Δ14^* + 2.5 µM Rapamycin (n = 18 embryos/54 neurons); ****p<0.0001. (N) Analysis of branch density at 60 hpf in *kptn^+/+^* (n = 17 embryos/51neurons), *kptn^Δ5/ Δ14^* (n = 17 embryos/51 neurons), *kptn^Δ5/ Δ14^* + 200 nM Rapamycin (n = 17 embryos/51 neurons) and *kptn^Δ5/ Δ14^* + 2.5 µM Rapamycin (n = 17 embryos/51 neurons); ****p<0.0001. Data represent pooled measurements from at least three independent biological replicates. Individual points correspond to single replicate in (B-D) and single neurons in (M,N); the mean is indicated by a dotted line, with error bars denoting ± SEM. Statistical significance was assessed using the Kruskal–Wallis test with Dunn’s multiple-comparison correction (M) and the Brown-Forsythe and Welch ANOVA test for experiments with more than two groups (N). Scale bar: 10 μm (E–H, I-L).

To directly evaluate the role of mTORC1 signalling in motor neuron hyperbranching in *kptn* mutants, we tested whether mTORC1 inhibition by rapamycin could rescue this phenotype. Zebrafish were evaluated at 26 or 60 hpf (similar to 3 and 5 dpf, no increase in p-RS6 was observed in *kptn* mutants in comparison with wild-type animals; Figure S8) following treatment with rapamycin. Chronic (200 nM added at 10 hpf and evaluated at 26 hpf, or added at 28 hpf and evaluated at 60 hpf) or acute (2.5 µM for 4 h before evaluation at 26 hpf or 60 hpf) treatments were used in these experiments. Both acute and chronic treatments could decrease mTORC1 activity, as indicated by reduced p-RS6 (Figure S8). At 26 hpf, *kptn^Δ5/Δ14^* treated with chronic (0.42 ± 0.02 protrusions/µm) or acute doses of rapamycin (0.39 ± 0.01 protrusions/µm) still displayed increased protrusion density compared to *kptn^+/+^* (0.27 ± 0.01 protrusions/µm) and were comparable to *kptn^Δ5/Δ14^*. Similar results were obtained at 60 hpf. The branch density of rapamycin-treated 60 hpf *kptn^Δ5/ Δ14^*, both chronic (0.14 ± 0.003 branches/µm) and acute (0.15 ± 0.004 branches/µm), remained significantly higher than *kptn^+/+^* (0.1 ± 0.003 branches/µm) (Figure 8E-N). These results reveal that the hyperbranching phenotype observed in the *kptn* mutants is independent of mTORC1 signalling. Taken together with earlier results, the regulation of axonal branching appears to be exclusively associated with Kptn’s actin-regulatory function.

## DISCUSSION

Interstitial collateral branching allows a single axon to contact multiple targets efficiently while preserving appropriate connectivity. Because the density and targeting of these collaterals directly control synaptic integration and homoeostasis, branch initiation must be tightly regulated; aberrant branching disrupts homoeostatic synaptic scaling and promotes network hyperconnectivity, whereas reduced branching compromises target innervation and causes developmental deficits. Positive regulators of axonal F-actin assembly, such as Ena/VASP proteins and Fmn2, license the initiation of collateral branches by promoting the maturation of axonal actin patches into filopodial protrusions. (Armijo-Weingart and Gallo, 2017; Dwivedy et al., 2007; Kundu et al., 2022; Nagar et al., 2021)However, the activities that oppose unconstrained branching have remained poorly defined. Here we show that Kptn, an F-actin barbed-end-binding factor with capping activity (Dutta et al., 2026), restricts axonal F-actin patch size and lifetime, thereby limiting the probability that a nascent patch matures into a protrusion and, ultimately, a stable collateral branch. Notably, Kptn depletion does not affect growth cone filopodia, indicating a local, compartment-specific function in neurons.

Axonal F-actin patches exist transiently, either dissipating or progressing to a protrusion depending on the regulatory inputs they receive, with increased patch size and persistence favouring productive, protrusion-competent patches (Gallo, 2024; Kundu et al., 2022). Viewed within this framework, our data place Kptn alongside other locally acting regulators of patch dynamics, including the Arp2/3 complex, Fmn2 and ADF/cofilin (Gallo, 2024; Kundu et al., 2022; Spillane et al., 2011; Tedeschi et al., 2019), as a critical component of the actin-regulatory network that determines whether a given site along the axon commits to branch formation, and identify barbed-end capping as a previously missing component of this network.

A central implication of this work is that axon collateral branching is governed by a quantitative balance between opposing barbed-end activities: Fmn2 promotes filament elongation and stabilises actin patches, increasing the likelihood of a patch-to-protrusion transition, whereas Kptn restricts filament elongation within the same structures. The inability of the actin-binding-deficient Kptn mutants to rescue F-actin patch and protrusion phenotypes *in vitro* and *in vivo* identifies F-actin barbed-end binding and restriction of actin filament elongation as a critical determinant of Kptn function. One attractive possibility is that Kptn and Fmn2 compete, directly or indirectly, for access to the barbed end within individual patches, such that their relative occupancy dictates the extent of filament elongation and, in turn, whether a given patch dissipates or initiates a protrusion. This model is supported by the epistatic interactions we observe: simultaneous depletion of both factors, or combined loss of their zebrafish orthologues, restores actin patch area, patch lifetime, protrusion density and branch density to wild-type levels, indicating that the two activities act on the same rate-limiting step of patch maturation. We suggest that this Kptn–Fmn2 axis functions as a local, tuneable mechanism setting the threshold for protrusion initiation along the axon shaft.

The consequences of dysregulated collateral branch initiation extend beyond axon morphology to the neuromuscular circuit and to behaviour (Koh et al., 2021; Ono et al., 2004). In *kptn* mutant zebrafish, excess CaP branching was accompanied by a proportional increase in neuromuscular junction number, whereas NMJ density per unit arbour length was unchanged. This suggests that Kptn does not overtly influence NMJ development, and that the increase in synapse number is principally a consequence of the expanded arbour providing more substrate for synaptogenesis. This aberrant, excess connectivity manifests as motor performance deficits, with *kptn* mutants displaying more frequent but lower-amplitude spontaneous tail coiling and an impaired touch-evoked escape response. The increased tail-coiling frequency is consistent with expanded innervation of the myotome, increasing the likelihood of eliciting a twitch. The reduced amplitude of tail coiling and the compromised swimming velocity and peak acceleration may reflect a loss of neuromuscular transmission synchrony and uncoordinated muscle fibre recruitment, resulting in impaired force generation. We cannot exclude additional contributions from altered muscle physiology or impaired neuromuscular transmission, but the overall pattern argues that precise regulation of arbour size is required for coordinated neuromuscular output.

Kptn is a component of the KICSTOR complex (Lupton et al., 2026; Teng et al., 2025; Wolfson et al., 2017), a negative regulator of mTORC1 signalling, and loss of Kptn has been reported to increase mTORC1 activity in brain tissue in mice (Levitin et al., 2023; Rawlins et al., 2026). However, we failed to observe any increase in mTORC1 signalling in *kptn* mutant larvae at any developmental stage relevant to this study. Tellingly, pharmacological inhibition of mTORC1 with rapamycin, at both acute and chronic dosing sufficient to reduce ribosomal S6 phosphorylation, failed to rescue the branching phenotype of *kptn* mutant motor neurons. Together with the inability of barbed-end-binding-deficient Kptn to rescue any F-actin patch or branching phenotypes, this indicates that axonal hyperbranching depends specifically on Kptn’s actin-regulatory function and is not a downstream consequence of altered mTORC1 signalling.

Macrocephaly is commonly associated with KPTN-related neurodevelopmental disorder (Baple et al., 2014; Levitin et al., 2023; Rawlins et al., 2026), yet we did not detect any increase in head size in *kptn* mutant larvae within the developmental window examined here. In affected individuals, head circumference is typically normal at birth, and macrocephaly emerges progressively over the first years of life (Rawlins et al., 2026). A comparable postnatal trajectory of brain overgrowth has been reported in Kptn-null mice (Levitin et al., 2023). The early larval stages assessed in the present study may therefore preclude the detection of analogous phenotypes in zebrafish. If Kptn regulates axonal branching in CNS neurons, it is conceivable that axonal hyperbranching contributes to the intellectual disability and increased seizure susceptibility observed in affected individuals (Baple et al., 2014; Biswas et al., 2025; Horn et al., 2022; Pajusalu et al., 2015; Rawlins et al., 2026; Thiffault et al., 2020; Ullah et al., 2022)

In summary, we identify Kptn as a barbed-end-binding actin regulator that acts locally within the axon shaft to restrain the maturation of axonal F-actin patches, in direct functional opposition to the elongation factor Fmn2 and independently of its role in mTORC1 regulation. Loss of this restraint results in more numerous but less effectively coordinated neuromuscular connections and, correspondingly, impaired motor output. These findings identify Kptn as a key regulator of axon interstitial branching and extend the F-actin patch model of collateral branch initiation from a permissive, elongation-driven process to one that is bidirectionally tuned by opposing barbed-end activities.

## MATERIALS AND METHODS

### Ethics approvals

All experimental protocols executed in this study received formal approval from the Institutional Animal Ethics Committee and the Institutional Biosafety Committee at IISER Pune.

### Zebrafish maintenance

All experiments were conducted using the wild-type TAB5 zebrafish strain. Adult breeding pairs were maintained at 28.5°C in recirculating aquaria (Techniplast) under a standard 14-hour light and 10-hour dark cycle. Breeding crosses were established overnight, and the resulting embryos were collected and raised at 28.5°C in E3 buffer (5 mM NaCl, 0.33 mM MgSO4, 0.17 mM KCl, 0.33 mM CaCl_2_, 5% Methylene Blue). These embryos were subsequently utilised at various developmental stages, as characterised by (Kimmel et al., 1995). The TAB5 strain also served as the genetic background for outcrossing with established mutant and transgenic lines. Specifically, the transgenic line Tg(mnx1:GFP) was employed, in which the motor neuron-specific *mnx1* promoter drives green fluorescent protein (GFP) expression. To inhibit melanin formation and ensure optical clarity for immunostaining and live-imaging assays, the E3 medium was supplemented with 0.003% Phenylthiourea (PTU; Sigma).

### Whole-mount In Situ hybridisation

Total RNA was isolated from 72 hpf TAB5 wildtype embryos using the RNeasy Mini Kit (Qiagen) and reverse-transcribed into cDNA utilising a reverse transcription kit (Takara). The resulting cDNA was used as a template to amplify a 331-bp fragment encompassing exon 12 and the 3’ untranslated region (UTR) of the zebrafish *kptn* gene. This amplicon was generated using primers specifically designed to flank the target sequence with T3 and T7 promoters in the sense and antisense directions, respectively (Kptn_3’UTR_T3: 5’-GCAATTAACCCTCACTAAAGGGCTGAAGACAACAGCGGATC-3’; Kptn_3’UTR_T7: 5’-TAATACGACTCACTATAGGGGACTATAACGATGACACTGATGC-3’). The amplified product was subsequently used for *in vitro* transcription to synthesise sense and antisense RNA probes targeting the *kptn* gene. Whole-mount *in situ* hybridisation was conducted on zebrafish embryos spanning developmental stages from the 1-cell stage to 72 hpf, following the protocol described by (Thisse and Thisse, 2008). Following staining, the samples were processed, stored in 80% glycerol (HiMedia), and subsequently imaged.

### Generation and Genotyping of *kptn* Knockout Zebrafish

Homozygous *kptn* mutants were generated using CRISPR Cas9-mediated genome editing. An sgRNA targeting exon 1 of the zebrafish *kptn* gene was designed using Benchling. Oligonucleotides containing the target sequence and an upstream T7 promoter were annealed and used as a template for *in vitro* transcription using the HiScribe T7 High Yield RNA Synthesis Kit (New England Biolabs). For microinjections, an injection mix containing 300 pg of Cas9 protein (Sigma CAS9PROT) and 50 pg of synthesised sgRNA, in a total volume of 2 nL, was injected directly into the cell of single-cell-stage embryos to generate F0 founders.

sgRNA sequence targeting *kptn* (Exon 1): 5’-GGCAGGCGAGCAGGAGTTGC-3’

Injected embryos were raised to sexual maturity and individually outcrossed to wild-type fish to identify founders carrying germline mutations. To screen the resulting F1 progeny, a 100 bp region flanking the PAM site was PCR-amplified and resolved on a 4% agarose gel to detect insertions or deletions. using the following primers: forward (5’-CTTCGCAGAGTAACATCTAC-3’) and reverse (5’-GATCTTCTTCTGTAGGTCTTGG-3’). F1 individuals exhibiting a band shift were subsequently validated via Sanger sequencing, a separate primer set was utilised: forward (5’-CGCCTGACAGTAAACTCTAGC-3’) and reverse (5’-CCAGTTCACTCCTTTCCAG-3’). Sequencing confirmed two independent deletions of 5 base pairs (bp) and 14 bp. F1 carriers of the 5 bp and 14 bp deletions were raised to adulthood and outcrossed to wild-type fish to dilute non-specific CRISPR-induced indels, and heterozygous mutants for each allele were confirmed by Sanger sequencing. Heterozygous siblings carrying the 5 bp or 14 bp deletion were then incrossed to generate the respective homozygous mutants, and the presence of stable homozygous lines was verified by Sanger sequencing (Figure S3).

### Site-directed mutagenesis and mRNA injection

Total RNA isolation and cDNA synthesis were performed as previously described. The entire *zkptn* cDNA was amplified using the forward primer 5’ GATTCGAATTCGCCACCATGTTACCCGACATAAAAGG 3’ and the reverse primer 5’ AGAGGCTCGAGCGTGTAGGGAAGCTTCAGT 3’. The resulting product was cloned into the pCS2-mNeonGreen vector using restriction digestion. Site-directed mutagenesis was performed following the protocol by (Liu and Naismith, 2008).The primers used for the mutagenesis were: Forward primer 5’ TACCAAGACCTACAGGATAAGATCAGACCTGTG 3’ and reverse primer 5’ CACAGGTCTGATCTTATCCTGTAGGTCTTGGTA 3’. The mutation was confirmed via Sanger sequencing. Capped mRNA was synthesised via *in vitro* transcription using the SP6 mMessage mMachine Kit (Invitrogen). This was performed from a linearised DNA template containing the SP6 promoter sequence upstream of the *zkptn^K55D^-*mNeonGreen construct. The mRNA was subsequently purified using the RNeasy MinElute Cleanup Kit (Qiagen). Finally, 50 pg of the purified *zkptn^K55D^-*mNeonGreen mRNA was injected into *kptn* mutants at the one-cell stage.

### Drug Treatment

Rapamycin (Invitrogen) was used to rescue mTORC1 hyperactivity and assess its effect on motor neuron branching. A 10mM stock solution was prepared and stored at −20°C for subsequent experiments. Both acute and chronic treatment regimens were performed via bath application in E3 buffer. For chronic treatment, at 26 hpf a concentration of 200 nM for 16 h was used and at 60 hpf the same concentration was used for 32 h. Whereas acute treatment utilised a 2.5 µM concentration for 4 h. For the 26 hpf branching rescue experiments, chronic doses were administered at 10 hpf, and acute doses at 22 hpf. For the 60 hpf experiments, chronic and acute treatments were initiated at 28 hpf and 56 hpf, respectively. For chronic treatment, rapamycin was replenished at 48 hpf. Following treatment, embryos were anaesthetised and mounted laterally in 1% low-melting-point agarose for imaging.

### Chick embryonic spinal neuronal culture

Primary culture of chick embryonic spinal cord tissue was done as mentioned in Kundu et al 2022. Embryonic spinal cord tissue from 6-days old *Gallus gallus* embryos were dissected out in dissecting medium [L-15 medium (Gibco, HiMedia) + 1X Penicillin-Streptomycin (HiMedia). Next, the extracted tissue was trypsinized with Trypsin - EDTA Solution 1X (0.25% Trypsin and 0.02% EDTA in Hank’s Balanced Salt Solution; HiMedia) at 37 °C for 20 minutes. For transfection, the cells were then resuspended in OptiMeM® (Gibco) media and electroporated using the NepaGene Super Electroporator NEPA21 Type II. The transfected tissue was then resuspended in culture medium [L-15 medium + 10% Fetal Bovine Serum (FBS, Gibco) + 1X Pen-Strep + 1X Sodium Pyruvate (HiMedia)] and plated on 75 µg/ml of Poly-d-lysine (Sigma-Aldrich) coated and 20 µg/ml Laminin (Sigma-Aldrich) coated glass-bottom 35mm plastic dishes. The cells were incubated with the culture medium at 37°C for 36–72 h.

### Transfection of spinal neuronal tissue with morpholinos and plasmids

The morpholinos used in this study are listed in Table S1. Similarly, the plasmids used are discussed in Table S2. All morpholinos were procured from Gene Tools, LLC. The knock-down efficiency of the morpholino targeted against the *Gallus gallus* Kptn (Kptn MO) has been verified, as shown in Figure S6. The morpholino against *Gallus gallus* Fmn2 (Fmn2 MO) has been extensively characterised for knock-down efficiency in previous studies (Ghate et al., 2020; Kundu et al., 2022; Sahasrabudhe et al., 2016). A standard control morpholino (Ctrl MO) was used as a negative control in all experiments. All transfections were performed together with a fluorescent reporter plasmid to identify successfully transfected cells. Only neurons exhibiting reporter fluorescence were selected for imaging and quantitative analysis.

### Fixation and immunofluorescence

#### Whole-mount immunostaining of zebrafish larvae

To label the neuromuscular junctions, embryos treated with PTU (Sigma) were collected at 60 hpf and stored in 4% paraformaldehyde (HiMedia) overnight at 4°C. Permeabilisation with 1mg/ml collagenase (Sigma) for 20 minutes was performed next. To mark the postsynapses, α-bungarotoxin conjugated with Alexa Fluor 555 (1:100; Invitrogen) was diluted in a blocking buffer containing 5% goat serum (HiMedia), 3% Bovine Serum Albumin (BSA; HiMedia), and 1% DMSO (HiMedia), and incubated for 2 h at room temperature. Next, to label the presynapses, the znp1 antibody (1:350; raised in mouse; DSHB) diluted in the aforementioned blocking buffer was used and incubated overnight at 4°C. This was followed by incubation with the Alexa-Fluor 488-conjugated anti mouse secondary antibody (1:1000; Invitrogen). For imaging, the embryos were then cleared in 50% glycerol.

#### For zebrafish larvae muscle staining

60 hpf embryos treated with PTU (Sigma) were collected stored in 4% paraformaldehyde (HiMedia) overnight at 4°C. Next, permeabilisation with 2% PBS-Triton X-100 (HiMedia) for 1.5 h at room temperature was done. Alexa-Fluor 488-conjugated Phalloidin (1:200; Invitrogen) diluted in a blocking buffer containing 5% goat serum, 3% BSA, and 1% DMSO was used to mark F-actin bundles of the myotome and incubated for 4 h at room temperature. For imaging, the embryos were then cleared in 50% glycerol.

#### For imaging neuronal morphology and axonal actin patches

After 36 h in vitro, neuronal cultures were fixed in 4% paraformaldehyde (Electron Microscopy Sciences) and 0.25% glutaraldehyde (Electron Microscopy Sciences) prepared in 1X PHEM buffer [60 mM PIPES (HiMedia), 25 mM HEPES (HiMedia), 10 mM EGTA (HiMedia), and 4 mM MgSO₄·7H₂O (HiMedia)]. Following fixation, cultures were permeabilised with 0.5% Triton X-100 (HiMedia) in 1X PHEM for 5 min at room temperature. Blocking was carried out using 3% BSA (HiMedia) in 1X PHEM (Blocking Buffer) for 1 h at room temperature. Primary anti-GFP antibody (raised in rabbit, Invitrogen) was applied at a dilution of 1:250 prepared in Blocking Buffer and incubated overnight at 4°C. Cultures were subsequently incubated for 2 h at room temperature with Alexa Fluor 488-conjugated anti-rabbit secondary antibody (1:1000; Invitrogen, for GFP) together with Alexa-Fluor 568-conjugated Phalloidin (1:100; Invitrogen, for F-actin) prepared in Blocking Buffer. Finally, mounting was done using Mounting Medium: 0.5% [w/v] n-propyl gallate (Sigma), 80% [v/v] glycerol, 20 mM Tris-HCl [pH 8.0], and 1× PBS.

#### For imaging microtubule innervation in axonal protrusions

Neuronal cultures were maintained for 72 h in vitro and subsequently fixed, permeabilized and blocked according to the previously described neuronal immunostaining protocol.

After blocking, cultures were incubated overnight at 4°C with primary antibodies diluted in blocking buffer: anti-βIII tubulin (1:3000; raised in mouse; Santa Cruz Biotechnology) and anti-GFP (1:250; raised in rabbit; Invitrogen). The following day, cultures were incubated for 2 h at room temperature with secondary antibodies diluted in blocking buffer: Alexa Fluor 568-conjugated anti-mouse antibody (1:2000; Invitrogen), Alexa Fluor 488-conjugated anti-rabbit antibody (1:1000; Invitrogen), and Phalloidin-647 (1:600; Invitrogen) for visualization of F-actin. The coverslips were mounted using n-propyl gallate-supplemented glycerol-based mounting medium.

### Image Acquisition

#### Zebrafish embryo and larvae morphology imaging

All whole mount *in situ* images for zebrafish larvae and the morphology at 3 dpf for wild type and *kptn* mutants were acquired on a Leica stereo zoom S8 APO microscope. For studying the morphology, larvae were first anaesthetised in 0.003% MS-222 (Sigma) and embedded in 1% low-melting agarose dorsally.

#### Zebrafish embryo mounting and imaging

All images were acquired on a Leica Microsystems TCS SP8 laser-scanning inverted confocal microscope using 20X/0.75 NA oil HC PL APO and 40X/1.30 NA oil HC PL APO objectives. Fixed embryos, previously stored in 50% glycerol, were mounted laterally onto 35 mm coverslip-bottom dishes using 1% low-melting agarose (Sigma). For live-cell imaging, embryos were first anaesthetised in 0.003% MS-222 (Sigma) and laterally embedded in 1% low-melting agarose to allow high-resolution imaging of the motor neurons. Image acquisition was performed using the Leica LAS X software.

#### Fixed neuronal imaging, including neuronal morphology, axonal actin patch analysis, and βIII-tubulin protrusion assays

Imaging was performed using a Leica Microsystems TCS SP8 laser-scanning inverted confocal microscope. For the 72 h βIII-tubulin protrusion experiments, the 63X/1.40 NA oil HC PL APO objective was used. Fixed imaging of neuronal morphology and axonal actin patch experiments was carried out using the 100X/1.4– 0.7 NA oil HCX PL APO objective. Image acquisition was performed using the Leica LAS X software.

#### Live imaging of F-actin patch dynamics

Neuronal cultures were transfected with the required morpholinos and/or plasmids and imaged between 36 and 48 h *in vitro*. Time-lapse imaging was performed using the Olympus (Evident) IX83 spinning disk confocal setup equipped with Hamamatsu Photonics ORCA-Flash 4.0 LT3 sCMOS camera (C11440) and a Tokai Hit stage-top incubator maintained at 37°C under humidified conditions. Images were acquired using a 100X/1.45 NA oil UPlanAPO objective and the Evident cellSens software. Only neurons exhibiting moderate F-tractin-mCherry expression were selected for imaging, as high expression levels of the actin-binding probe can induce excessive F-actin bundling or stabilization. Images were collected at 4 s intervals for a total duration of 5 min with 2 × 2 binning. Exposure times for the F-tractin-mCherry channel generally ranged from 200 to 600 ms, depending on signal intensity. The Z-drift compensation system (ZDC) was enabled throughout image acquisition.

#### Dual-channel live imaging of Kptn^WT^-GFP and F-actin patches

Neurons were transfected with pCAG-Kptn^WT^-GFP and pCAG-F-tractin-mCherry and time-lapse imaging was done as previously explained. Sequential acquisition of the two channels was performed, with images collected at 4 s intervals for a total duration of 5 min. As the exposure time of the first channel (F-tractin-mCherry) ranged from 200 to 600 ms, the effective temporal delay between the two channels was within the same range. ZDC was enabled throughout image acquisition.

### Image Analysis

All image analysis was carried out using Fiji (Schindelin et al., 2012).

For *in vivo* quantification of protrusion density and branch density at 26 hpf and 60 hpf, respectively, of zebrafish motor neurons, the length of the CaP neuron at those stages was considered to normalise the total number of protrusions and total number of branches to get protrusion density and branch density. The number of protrusions, branches and length of CaP neuron was calculated manually using the ‘Segmented line’ tool.

For all *in vitro* analyses, a minimum axonal length threshold of 50 µm was applied to distinguish axons from shorter neurites. Both axonal and growth cone protrusions were traced manually using the ‘Segmented line’ tool. For analysis of axonal protrusions, only structures measuring at least 0.5 µm were traced manually, whereas a minimum cut-off of 2 µm was used for growth cone filopodia. Total axon length was also measured manually. Axonal protrusion density was then calculated as the number of protrusions per unit axon length (number of protrusions/total axonal length). To determine growth cone area, images were first thresholded using the ‘Adjust Threshold’ function in Fiji to generate a binary mask. The growth cone region was then selected from the mask using the ‘Wand’ tool, and the enclosed area was recorded. To quantify βIII-tubulin invasion into axonal protrusions, all Phalloidin-labelled protrusions exceeding 0.5 µm in length were identified, manually traced, and measured. Individual protrusions were then assessed for the presence of βIII-tubulin signal, and those containing detectable βIII-tubulin were classified as βIII-positive. βIII-positive protrusion density was then calculated by normalising the number of βIII-positive protrusions to the corresponding axon length.

For analysing axonal actin-patch area from fixed images, the patches were manually selected and their area and density (number of patches per unit length of axon) were calculated.

For analysis of actin patch dynamics from time-lapse images, image sequences were first corrected for photobleaching using the ‘Histogram Matching’ method (Miura, 2020). Correction of lateral (x–y) drift was performed using the ‘Fast4DReg’ plugin (Laine et al., 2019; Pylvänäinen et al., 2023). A segmented line was drawn manually along the longitudinal axis of the axon, and kymographs were generated using the ‘KymoToolBox’ plugin (Zala et al., 2013). Individual actin patch trajectories were manually traced on the kymographs using the ‘Segmented line’ tool and saved as ROIs. These ROIs were analysed using the ‘Analyse Kymo’ function of ‘KymoToolBox’ to obtain patch lifetime. Within the plugin, movement was defined as velocities exceeding 0.1 µm/s.

### Western Blotting

#### Lysate preparation and immunoblotting for probing mTORC1 activity in zebrafish

Larvae at 3, 5- and 7-days post-fertilisation (dpf) were anaesthetised and deyolked, followed by lysate preparation in RIPA buffer (50 mM Tris-HCl (pH 7.4-8.0), 150 mM NaCl (HiMedia), Triton X-100, 0.5% sodium deoxycholate (HiMedia), 0.1% Sodium Dodecyl Sulphate (SDS; HiMedia), and 1-5 mM EDTA (Sigma) supplemented with protease inhibitor (Protease Inhibitor Cocktail; Roche, Phenylmethylsulfonyl Fluoride; Sigma) and phosphatase inhibitor (Cell Signalling Technology). Similarly, larvae treated with both chronic and acute doses of rapamycin at 26 hpf and 60 hpf were anaesthetised and deyolked and lysates were prepared in RIPA buffer. For homogenization, the samples were sonicated. The lysates were then centrifuged at 14,000 rpm for 20 minutes at 4°C. The supernatant was collected, and protein concentration was quantified using a BCA assay (Takara). For each sample, 20 μg of protein was loaded onto a 10% SDS-polyacrylamide gel. Following electrophoresis, proteins were blotted onto a PVDF membrane (Immobilon, Millipore Sigma). The membranes were blocked in 5% BSA dissolved in 0.1% PBS-Tween (PBST). Subsequently, immunoblotting was performed using primary antibodies against ꞵ-actin (1:10 000; raised in mouse; Santa Cruz Biotechnology), total S6 (1:7000; raised in rabbit; Cell Signalling Technology), phospho-S6 (1:7000; raised in rabbit; Cell Signalling Technology), GAPDH (1:5000; raised in rabbit; Sigma). Proteins were detected using HRP-conjugated anti-mouse (1:10,000; Invitrogen) and anti-rabbit secondary antibodies (1:10,000; Invitrogen). Finally, protein bands were visualised using ECL medium (Immobilon, Millipore Sigma) and imaged using a ChemiDoc iBright1500 (Invitrogen) imaging system.

#### Lysate preparation and immunoblotting for Kptn morpholino efficiency examination

Neuronal cultures were treated with either Ctrl MO or Kptn MO. For protein extraction, cultures were incubated till 36h and chilled 1× Laemmli buffer [5X Laemmli buffer: 250 mM Tris-HCl (pH 6.8), 30% (v/v) glycerol, 4% (w/v) SDS, 0.06% (w/v) bromophenol blue (HiMedia), and 16% β-mercaptoethanol (HiMedia)] was added directly to each culture dish. Cells were then scraped using a pipette tip. Lysates were collected into pre-chilled 1.5 ml microcentrifuge tubes. The samples were then boiled at 99°C prior to standard SDS-PAGE and western blot analysis.

The primary antibody against Kptn was a kind gift from Dr. Sankar Maiti, IISER Kolkata. The antibody was raised in mouse against amino acids 1–258 of the human Kptn protein as the epitope. The 7th bleed of the sera was used as the primary antibody at a dilution of 1:1000. HRP-conjugated anti-mouse secondary antibody (Invitrogen) was used at a dilution of 1:2500. GAPDH served as the loading control (primary antibody –1:3000, Santa Cruz Biotechnology; secondary antibody – HRP-conjugated anti-mouse, 1:10,000). To ensure equal protein loading between treatments, an initial GAPDH immunoblot was performed for each biological replicate and band intensities were quantified by densitometry to determine normalized sample loading volumes. Subsequently, samples were re-probed for Kptn immunoblotting.

### Behavioural experiments and setups

#### Spontaneous Tail Coiling (STC) Assay

Spontaneous tail coiling behaviour was assessed using embryos at 24 hpf. Embryos within their chorions were transferred to 35 mm Petri dishes containing E3 buffer maintained at 28.5°C. Tail coiling activity was recorded for 200 seconds at 15 frames per second (fps) using a digital video camera (AVT Pike, F-032B). Video analysis was subsequently performed using the previously published MATLAB script, ZebraSTM (González-Fraga et al., 2019).

Touch-Evoked Escape Response (TEER) Assay

Larvae at 60 hpf were placed in a 35 mm Petri dish and subjected to a tactile stimulus. The stimulus was delivered using a modified tuberculin needle with a soft nylon fibre attached to its tip. A gentle touch was applied to the tail of each larva, and the resulting escape behaviour was recorded using a high-speed video camera (AVT Pike, F-032B). The movement trajectories were analysed using the ‘Manual Tracking’ plugin in Fiji. From the resulting coordinates, the total distance travelled, average velocity, and maximum acceleration were calculated manually.

### Graphical representation and statistical analysis

All the graphical representation and statistical analyses were performed using GraphPad Prism 9.5. Graphical representations were plotted using scatter dot plot with the dotted middle line denoting the mean and error bars indicating the standard error of mean (SEM). Number of data points quantified in each graph have been indicated in the figure legends. Mann-Whitney *U* test and Welch’s t test was employed for comparing two groups. When comparing more than two groups, non-parametric one-way ANOVA or Kruskal-Wallis test was used and parametric Brown-Forsythe test was used. Two-way ANOVA test was used to compare groups across conditions.

## AUTHOR CONTRIBUTIONS

Conceptualisation: R.B., S.M. and A.G.; Investigation and formal analysis: R.B. and S.M.; Methodology and resources: R.B., S.M. and A.G.; Writing – original draft: R.B.,

S.M. and A.G.; Writing – review and editing: R.B., S.M. and A.G.; Funding Acquisition: A.G.

All authors gave final approval for publication and agreed to be held accountable for the work performed therein.

## ACKNOWLEDGEMENTS

The IISER Pune Microscopy Facility and the National Facility for Gene Function in Health and Disease (NFGFHD) at IISER Pune are acknowledged for access to imaging systems and maintenance of the zebrafish facility. We thank Dr. Sankar Maiti, IISER Kolkata, for the gift of the anti-Kaptin antibody, and Dr. Priyanka Dutta, Brandeis University, for providing human Kaptin cDNA.

## AI USE DECLARATION

The authors used standard AI tools to assist with language editing and improving readability. All content has been reviewed and edited by the authors, who take full responsibility for the accuracy and originality of the work.

## DATA AVAILABILITY

All data is available on reasonable request.

---

**Supplementary Figure 1.**
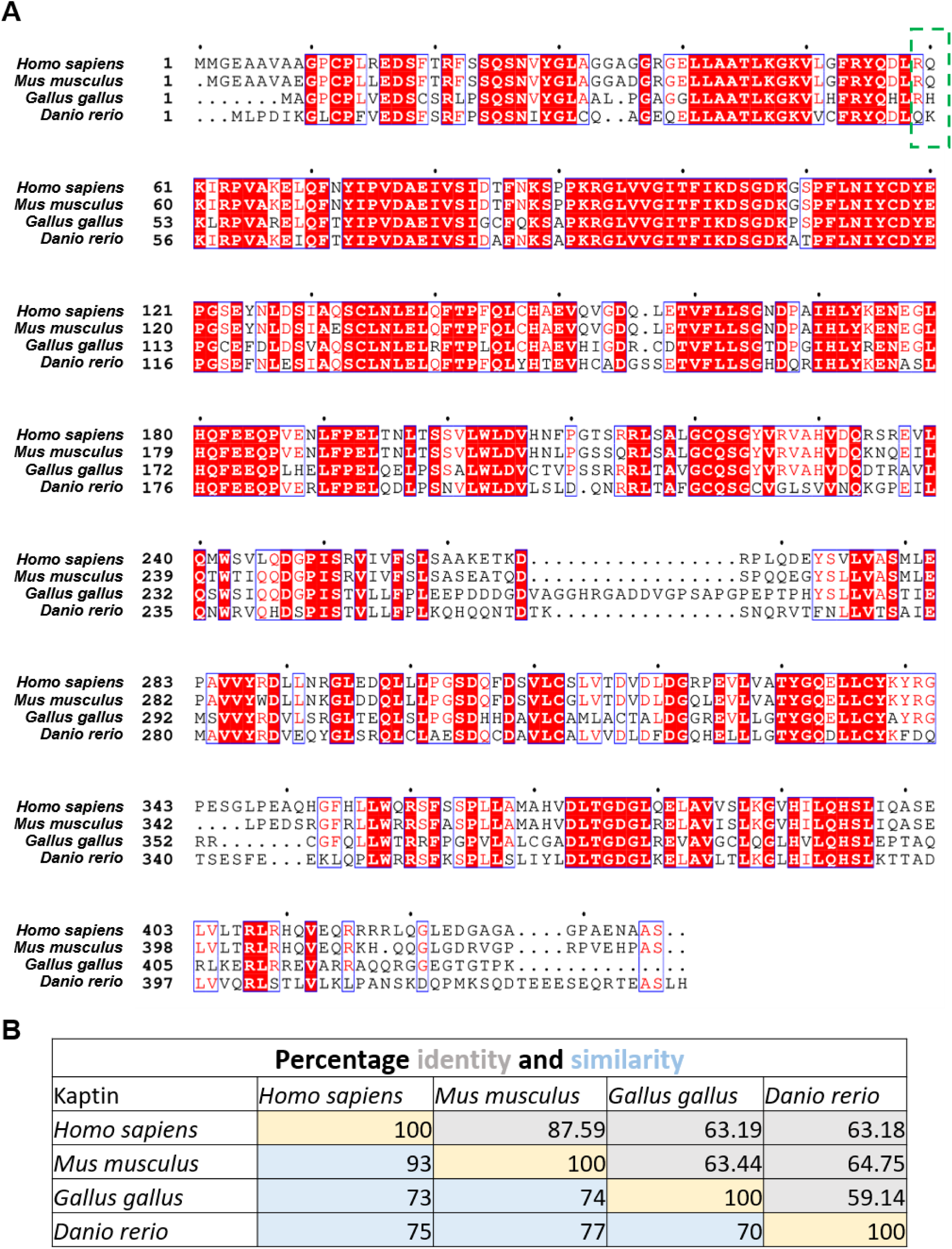
Multiple sequence alignment of Kptn across species. (A) Multiple sequence alignment of Kptn homologues from *Homo sapiens* (Uniprot ID-Q9Y664), *Mus musculus* (Q8VCX6), *Gallus gallus* (A0A1D5PJB7) and *Danio rerio* (Q6DC73). Sequences were aligned using Clustal Omega (Sievers and Higgins, 2014) and the alignment rendered with ESPript 3.2 (Gouet et al., 1999). Strictly conserved residues (identical across all sequences) are shown as white letters on a red background. Residues conserved in physicochemical character but not identity are shown as red letters and enclosed in blue boxes denoting similarity groups. Non-conserved residues are shown as black letters. Numbering on the left indicates residue positions for each sequence. The green dashed box highlights the actin-binding site of arginine at position 59 of *Homo sapiens* homologue. The equivalent position is lysine at 55^th^ position for *Danio rerio* homologue. (B) Table showing pairwise percentage identity and similarity of the Kptn protein across four vertebrate species. Amino acid sequences from *Homo sapiens, Mus musculus, Gallus gallus and Danio rerio* were aligned, and values are expressed as percentage identity (upper right) and percentage similarity (lower left) relative to the human reference sequence. Identity denotes positions with identical residues, whereas similarity additionally includes conservative amino acid substitutions. Sequences were retrieved from UniProt. Higher values indicate greater evolutionary conservation of the protein between species.

**Supplementary Figure 2.**
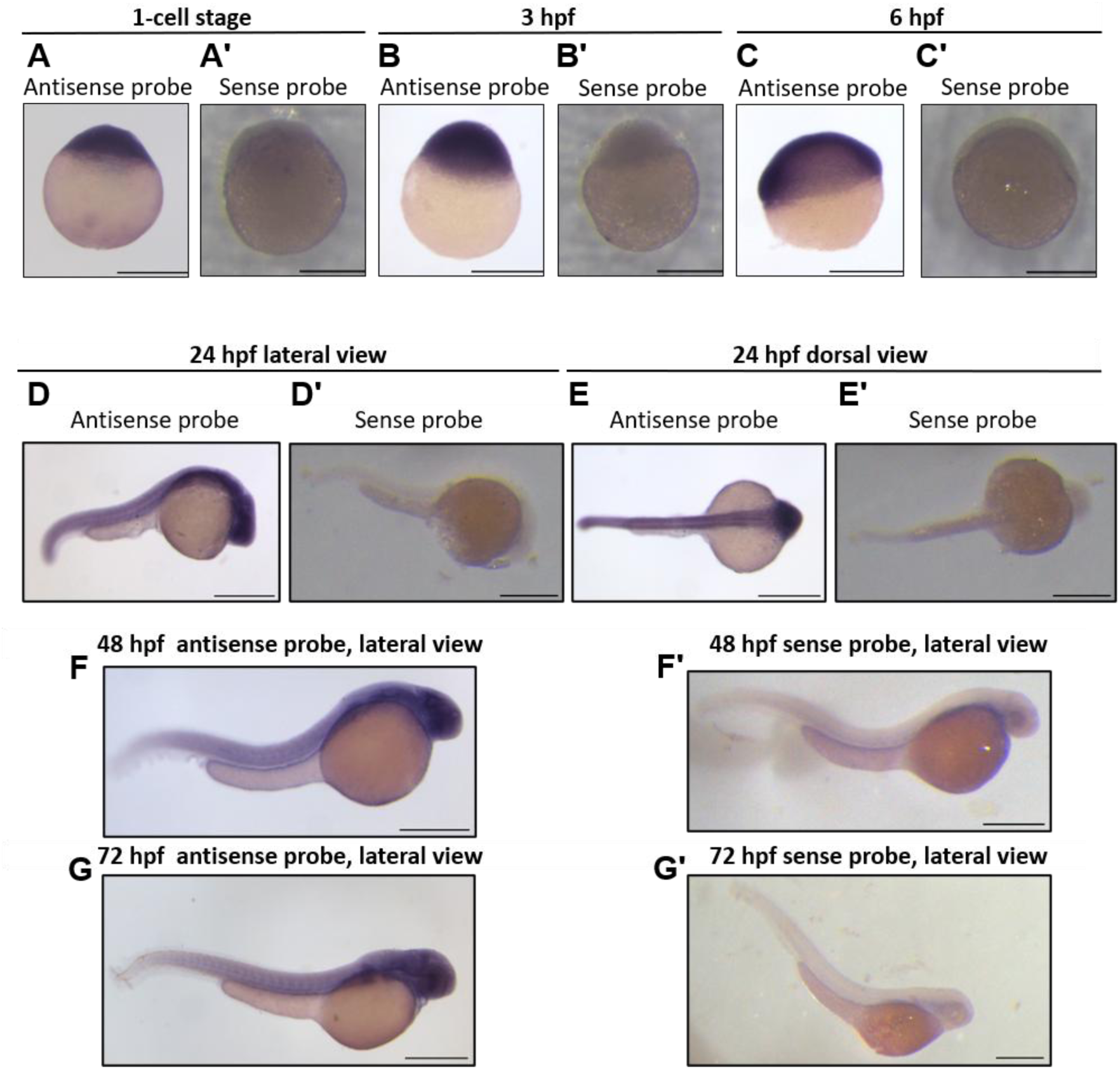
*kptn* is enriched in the zebrafish nervous system. (A-G) Representative images of *kptn* mRNA expression across various stages of development. (A) 1-cell stage antisense probe, (A’) 1-cell stage sense probe, (B) 3 hpf antisense probe, (B’) 3 hpf sense probe, (C) 6 hpf antisense probe, (C’) 6 hpf sense probe, (D) 24 hpf antisense probe, lateral view, (D’) 24 hpf sense probe, lateral view, (E) 24 hpf antisense probe, dorsal view, (E’) 24hpf sense probe, dorsal view, (F) 48 hpf antisense probe, lateral view, (F’) 48 hpf sense probe, lateral view, (G) 72 hpf antisense probe, lateral view, (G’) sense probe, lateral view. Scale bar: 500 μm (A-G).

**Supplementary Figure 3.**
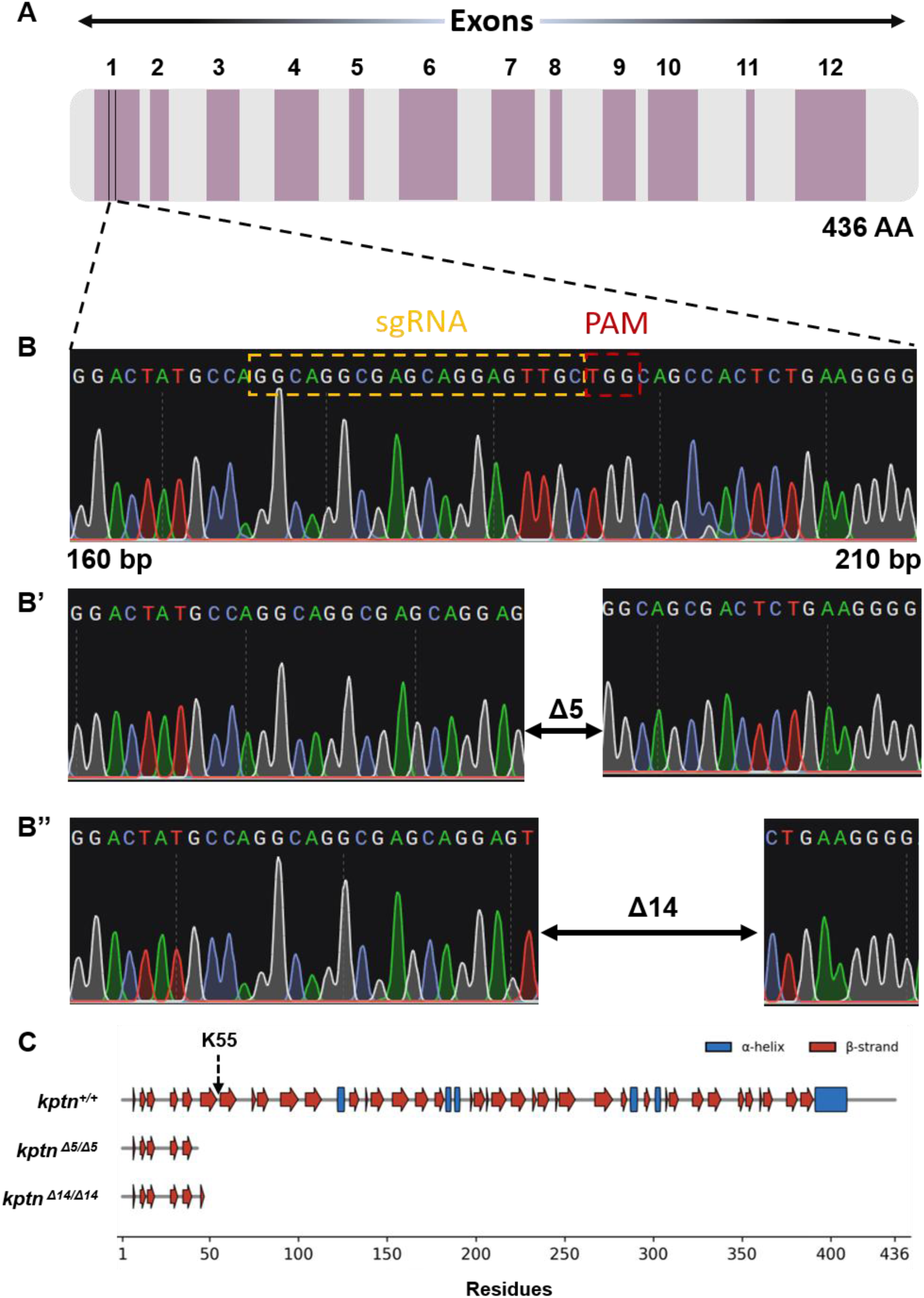
Generation of Zebrafish *kptn* mutant using CRISPR Cas 9. (A) Schematic depicting the full-length zebrafish *kptn* gene with 12 exons encoding for 433 amino acid. It marks (in black) the genomic location where sgRNA for *kptn* is designed in exon 1. B) Representative chromatogram for part of the wild type *kptn* amplicon (160 bp – 210 bp) sequenced by sanger sequencing. The yellow dotted line represents the sgRNA sequence followed by the PAM sequence in red. (B’) Representative chromatogram from homozygous CRISPR mutant showing a 5bp deletion. (B”) Representative chromatogram from homozygous CRISPR mutant showing a 14 bp deletion. (C) Secondary structure of wild-type Kptn and its two truncation mutants, assigned by DSSP from the AlphaFold-predicted model (UniProt Q6DC73) and mapped along the 436-residue sequence: α-helices (blue blocks), β-strands (red arrows), and coil (grey line). The actin binding site K at the 55^th^ position is marked with black dashed arrow. *kptn ^Δ5/Δ5^* and *kptn ^Δ14/Δ14^* carry premature stop codons, so they are drawn only to residues 43 and 46 respectively. This leads to the loss of nearly all downstream structure that is directly apparent on the shared residue axis.

**Supplementary Figure 4.**
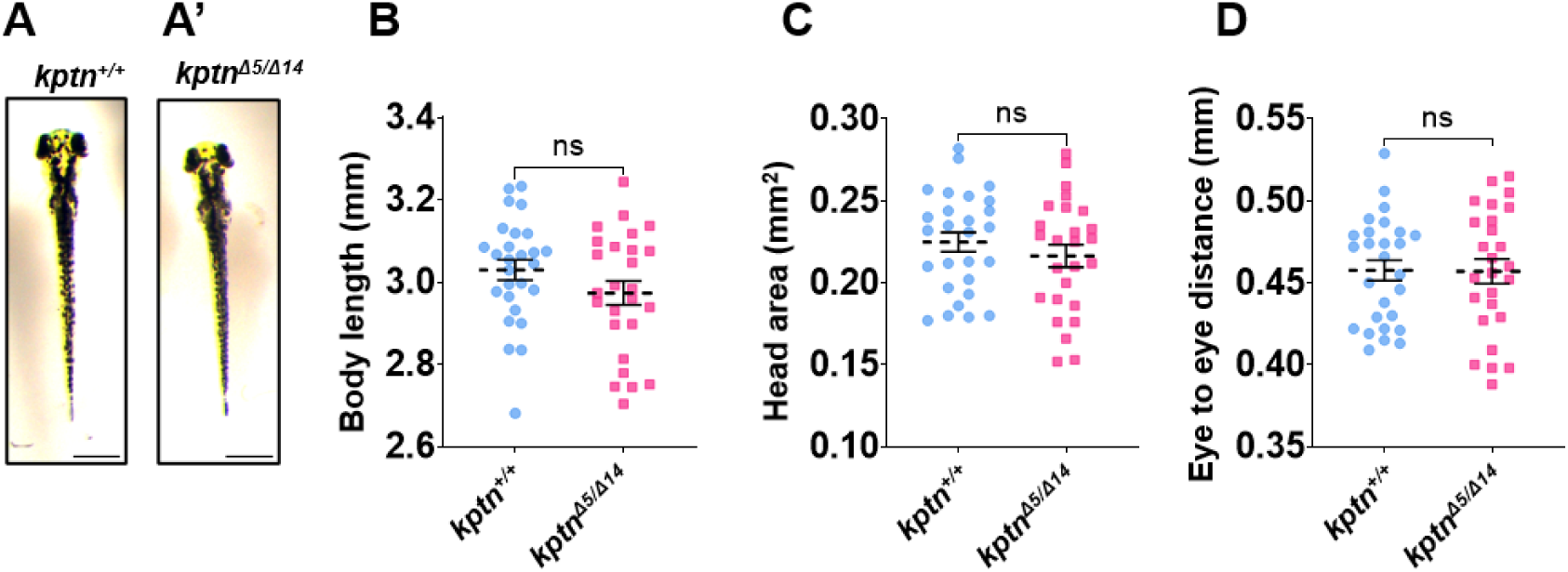
Morphological parameters are comparable between *kptn* mutants and wild type zebrafish. (A) Representative images of 3dpf *kptn^+/+^.* (A’) Representative images of 3 dpf and *kptn^Δ5/ Δ14^* larvae. (B) Quantification of body length at 3 dpf of *kptn^+/+^* (n = 27 larvae), and *kptn^Δ5/ Δ14^* (n = 26 larvae). (C) Quantification of head area at 3 dpf of *kptn^+/+^* (n = 27 larvae), and *kptn^Δ5/ Δ14^* (n = 26 larvae). (D) Quantification of eye to eye distance at 3 dpf of *kptn^+/+^* (n = 27 larvae), and *kptn^Δ5/ Δ14^* (n = 26 larvae); ns, p=0.1482; p=0.3581; p=0.9551. Data represent pooled measurements from at least three independent biological replicates. Individual points correspond to single embryo in (B-D); the mean is indicated by a dotted line, with error bars denoting ± SEM. Statistical significance was assessed using the Welch’s t-test for two-group comparisons (B-D). Scale bar: 500 μm (A and A’).

**Supplementary Figure 5.**
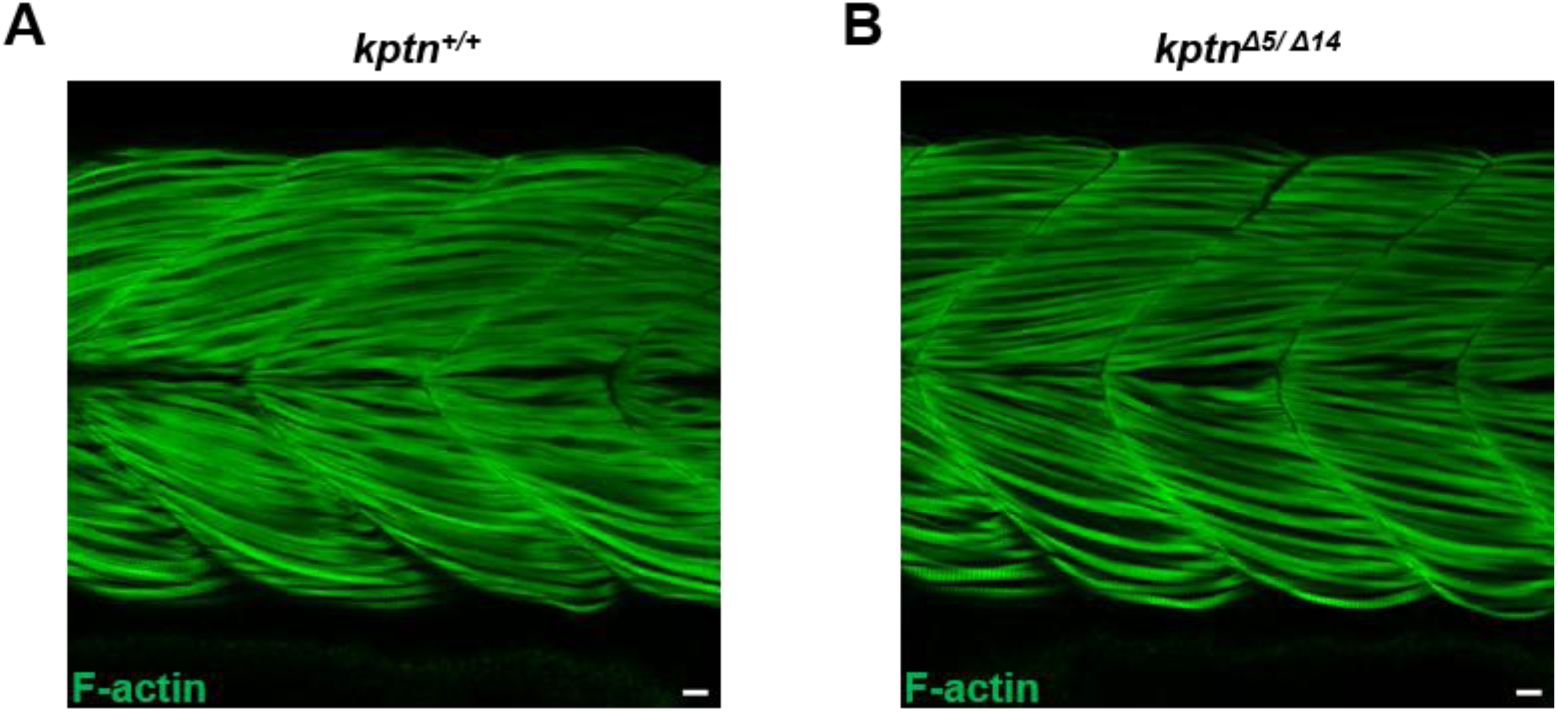
Muscle architecture is not affected in *kptn* mutants. (A and B) Representative micrographs of 60 hpf (A) *kptn^+/+^* (B) *kptn^Δ5/ Δ14^* probed with Alexa Fluor 488–conjugated to label F-actin bundles in the myotome of zebrafish. Visually, gross morphology of the muscles for *kptn* mutants looks similar to wild type. For both images, scale bar: 10 µm.

**Supplementary Figure 6.**
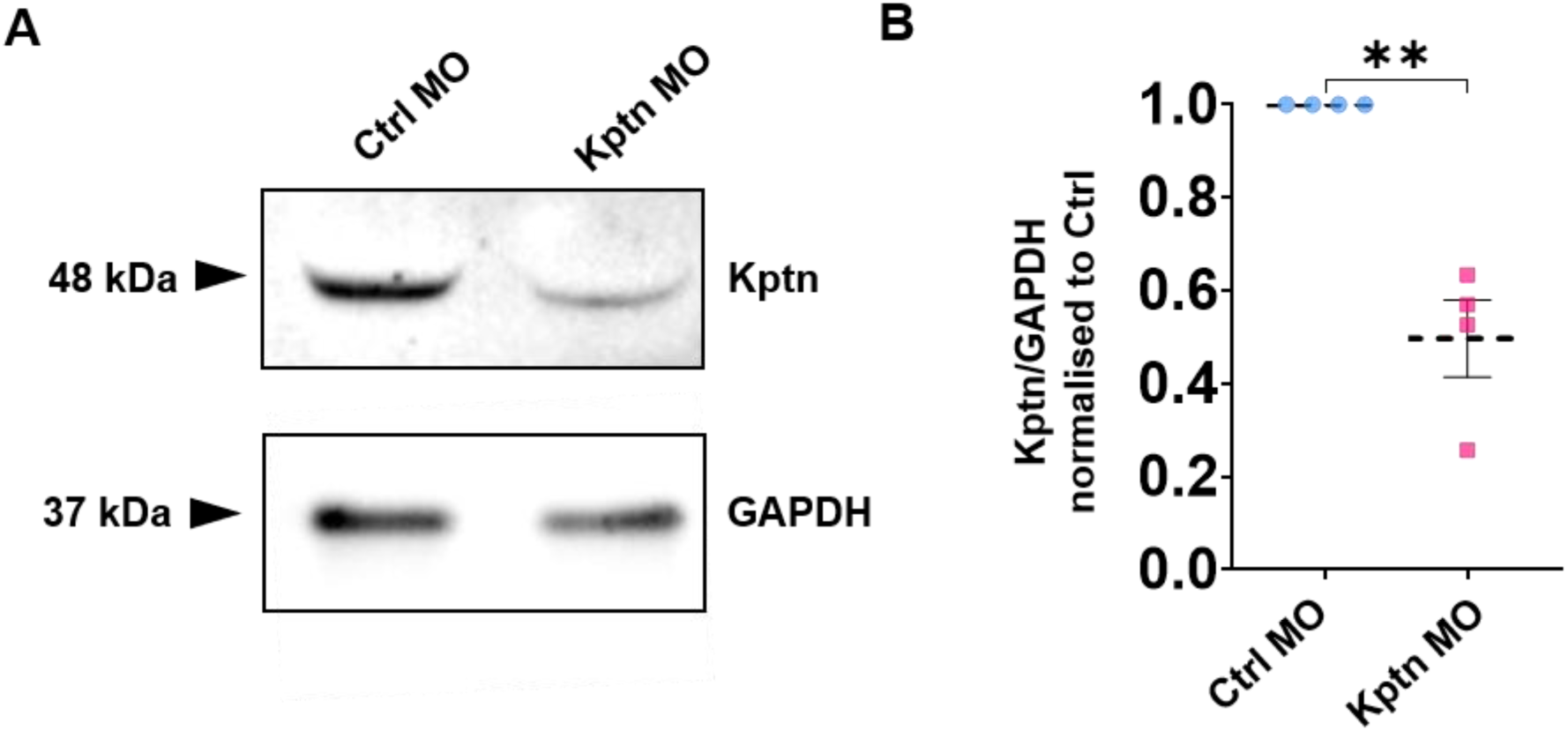
Verification of Kptn MO knock-down efficiency. (A) Representative immunoblot showing knockdown efficiency of translation blocking morpholino against Kptn (Kptn MO). B shows the quantification of Kptn protein levels by immunoblotting from neuronal lysates upon treatment with indicated morpholinos (**p=0.009). Data represent pooled measurements from at least three independent biological replicates. The mean is indicated by a dotted line, with error bars denoting ± SEM. Statistical significance was assessed using the unpaired t-test with Welch’s correction. Transfection of neurons with Kptn MO resulted in almost 50% reduction of Kptn protein levels (0.48 ± 0.08) relative to neurons transfected with the non-specific control morpholino (Ctrl MO). As the transfection efficiency achieved with our protocol is routinely under 40%, the protein reduction observed in immunoblots significantly underrepresents the knockdown efficiency. To specifically identify morpholino-transfected neurons for analysis, morpholinos were co-transfected along with a fluorescent reporter (GFP) tagged plasmid, and only reporter-positive neurons were selected.

**Supplementary Figure 7.**
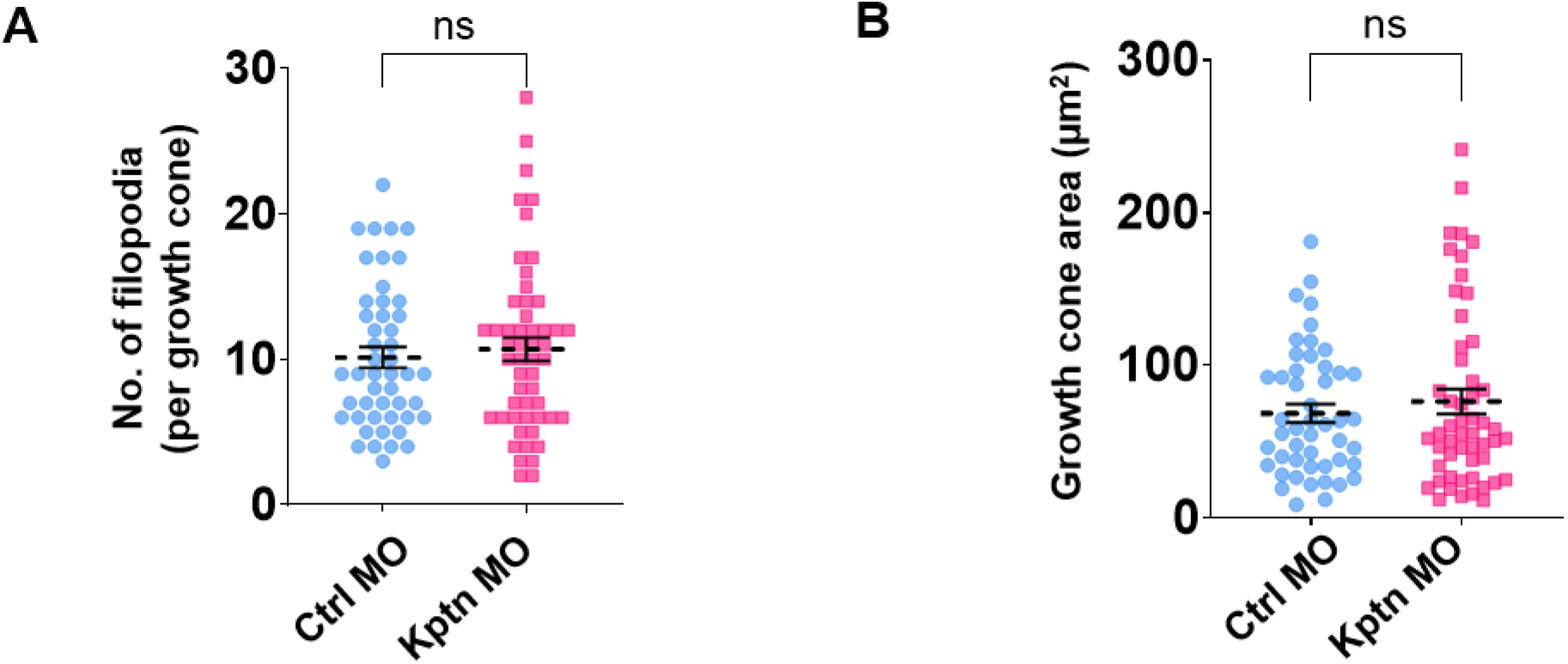
Kptn depletion does not affect growth cone filopodia. (A, B) Analysis of growth cone filopodial parameters from micrographs. A depicts the quantification of number of filopodia per growth cone in neurons treated by either Ctrl MO (n = 48) or Kptn MO (n = 53, ns, p=0.7458). D shows the analysis of growth cone area in Ctrl MO (n = 48) and Kptn MO treated (n = 53) neurons (ns, p=0.9649). Data are pooled from at least three independent biological replicates. The mean is indicated by a dotted line, with error bars denoting ± SEM. Statistical significance was assessed using the Mann–Whitney *U* test for two-group comparisons.

**Supplementary Figure 8.**
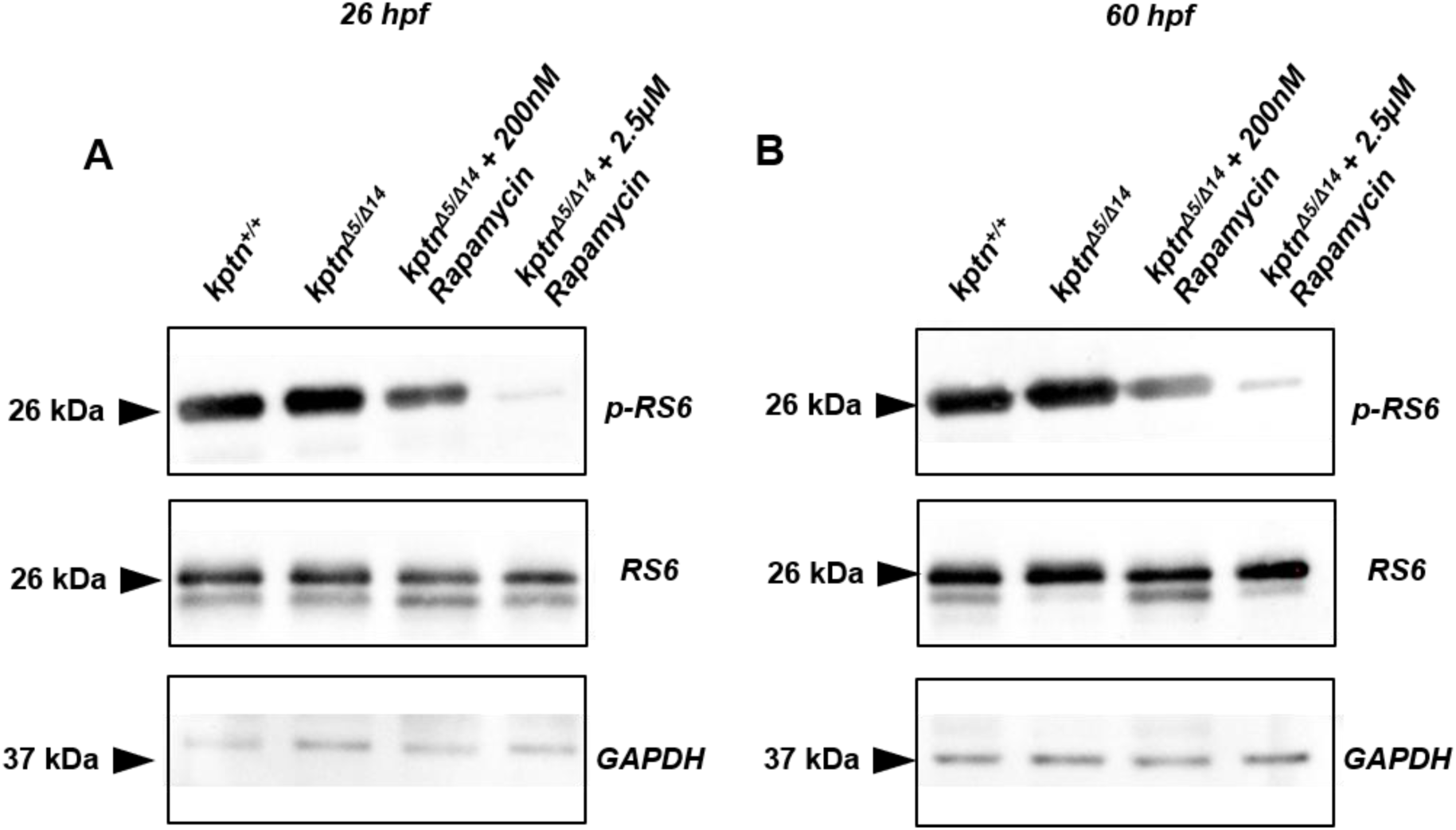
Rapamycin inhibits mTORC1 activity in *kptn* mutants. (A) Representative immunoblot for p-RS6 and RS6 protein in 26 hpf larvae for acute and chronic treatment of rapamycin. (B) Representative immunoblot for p-RS6 and RS6 protein in 60 hpf larvae for acute and chronic treatment of rapamycin.

**Table S1:**
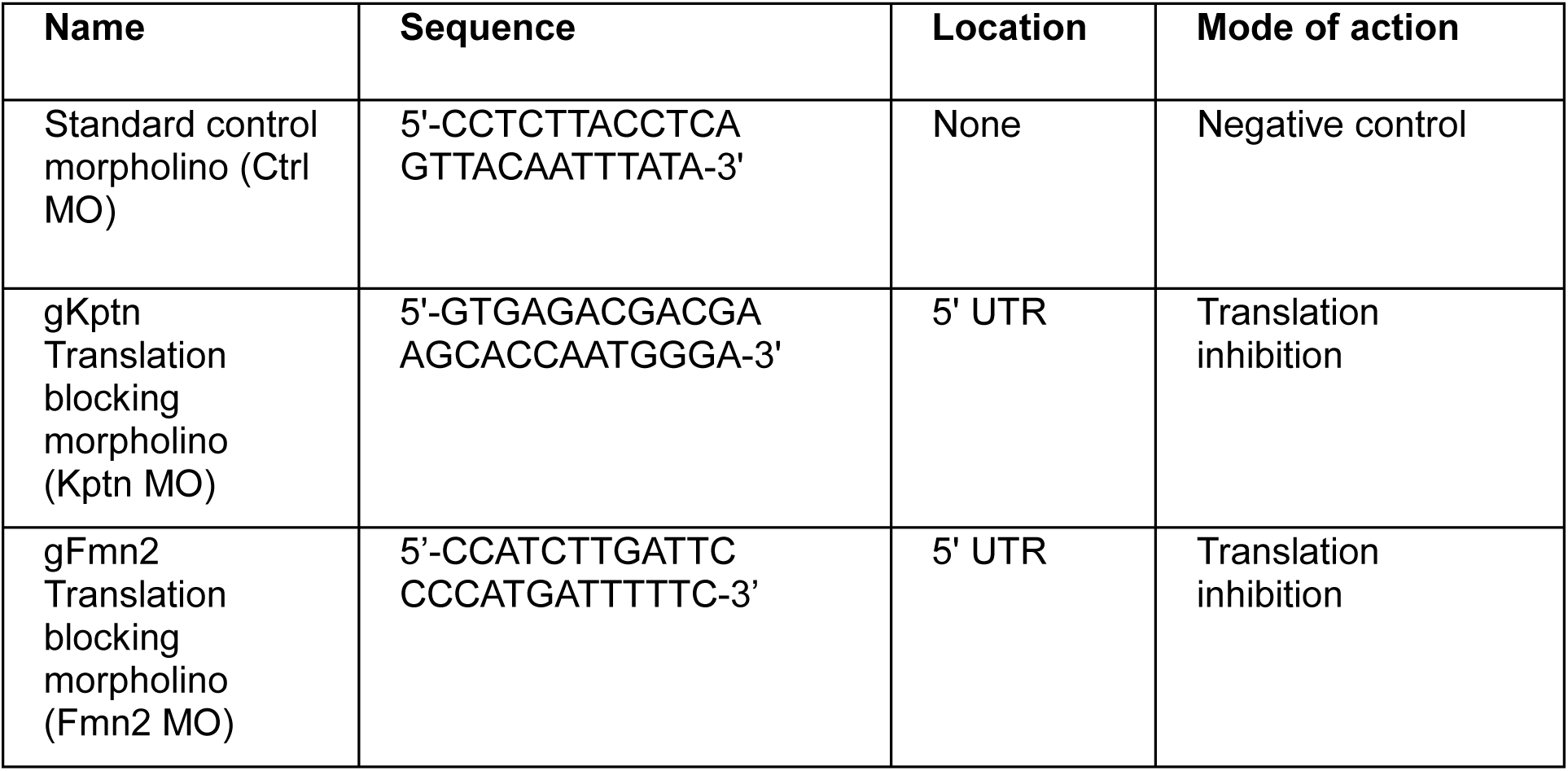
Morpholinos used in this study.

**Table S2:**
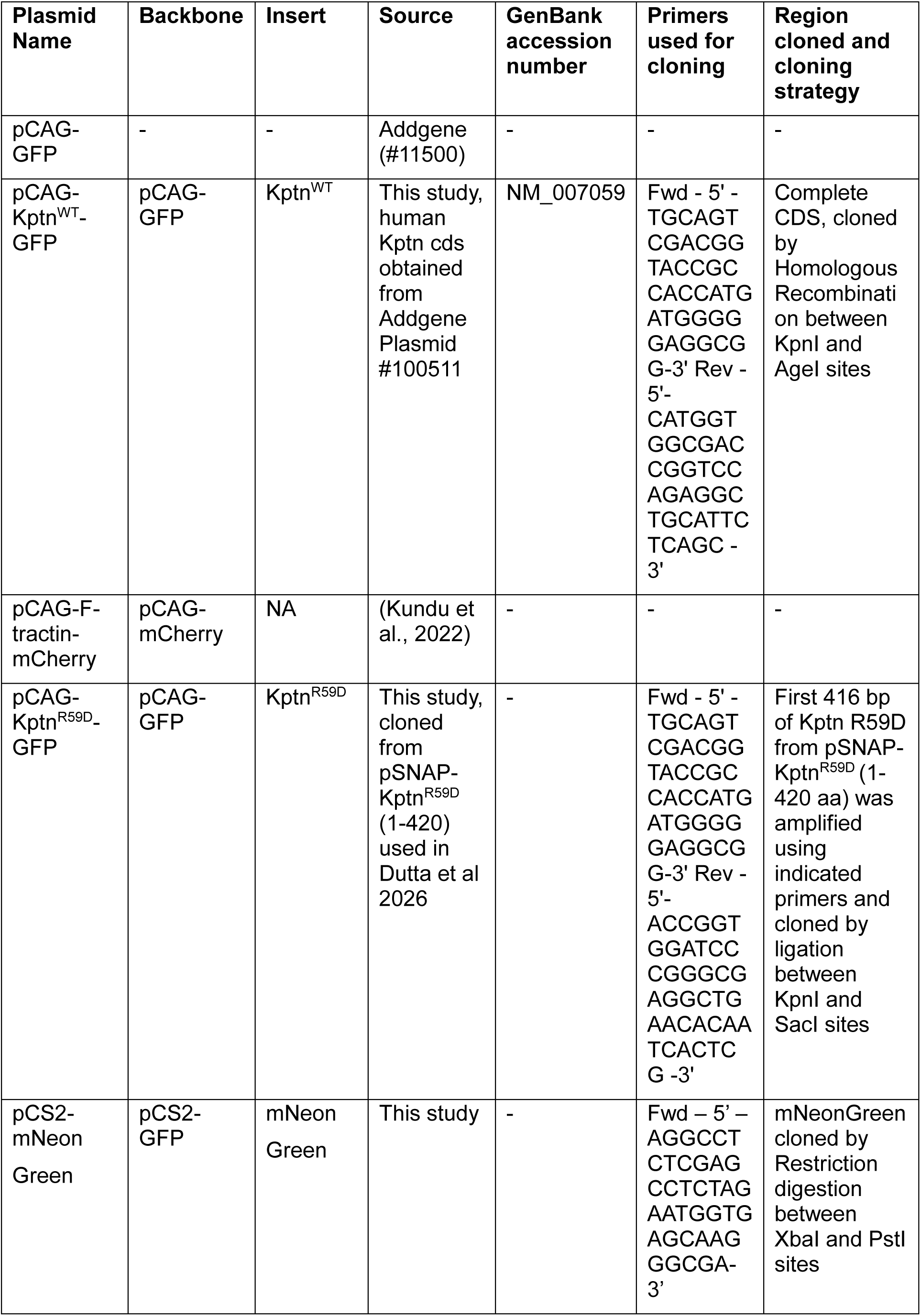
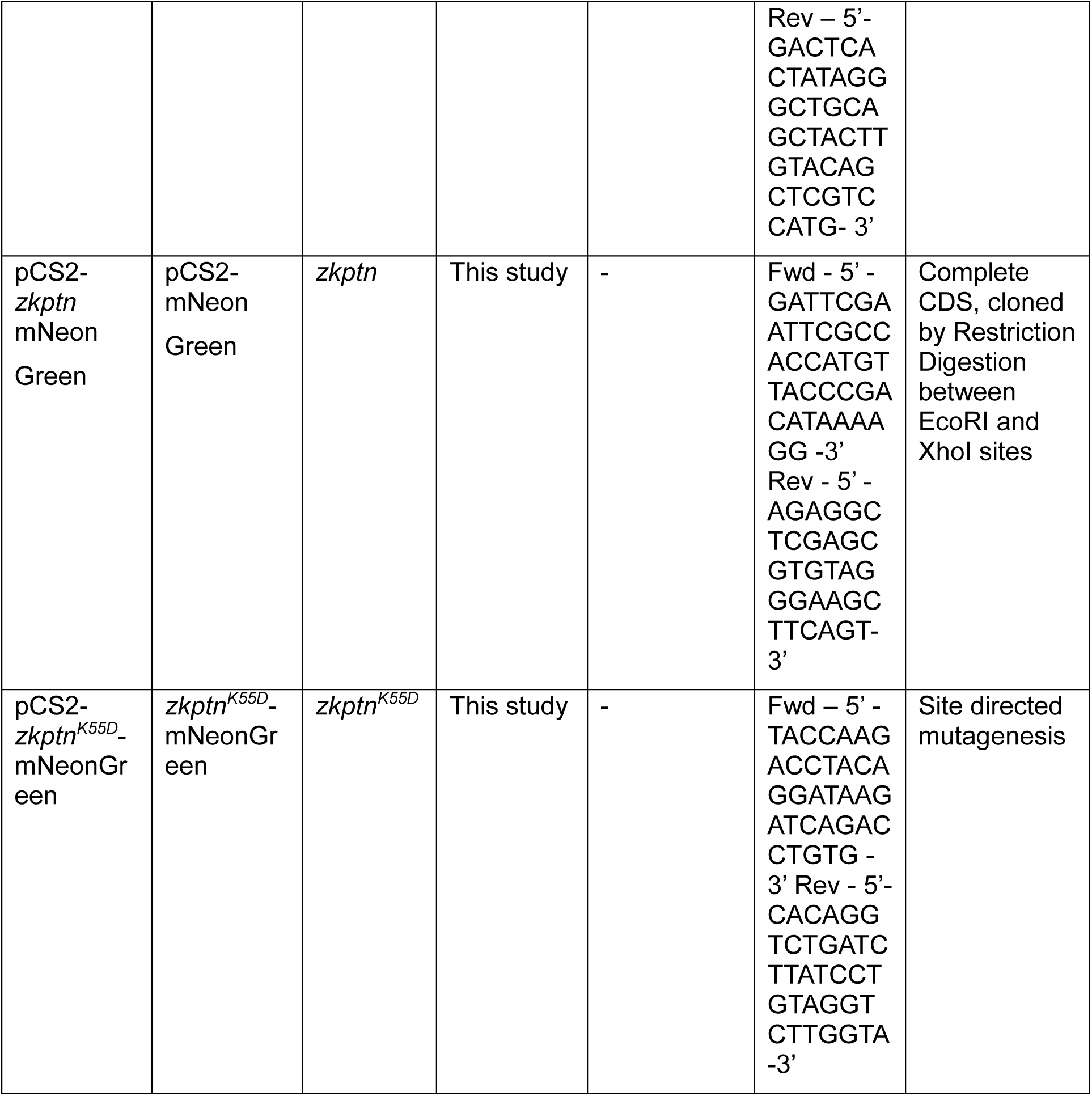
Plasmid constructs used in this study.

